# Convergent motifs of early olfactory processing are recapitulated by layer-wise efficient coding

**DOI:** 10.1101/2025.09.03.673748

**Authors:** Juan Carlos Fernández del Castillo, Farhad Pashakhanloo, Venkatesh N. Murthy, Jacob A. Zavatone-Veth

## Abstract

The architecture of early olfactory processing is a striking example of convergent evolution. Typically, a panel of broadly tuned receptors is selectively expressed in sensory neurons (each neuron expressing only one receptor), and each glomerulus receives projections from just one neuron type. Taken together, these three motifs—broad receptors, selective expression, and glomerular convergence—constitute “canonical olfaction,” since a number of model organisms including mice and flies exhibit these features. The emergence of this distinctive architecture across evolutionary lineages suggests that it may be optimized for information processing, an idea known as efficient coding. In this work, we show that by maximizing mutual information one layer at a time, efficient coding recovers several features of canonical olfactory processing under realistic biophysical assumptions. We also explore the settings in which noncanonical olfaction may be advantageous. Along the way, we make several predictions relating olfactory circuits to features of receptor families and the olfactory environment.

## I. INTRODUCTION

Chemosensation is our oldest sensory modality, and for most animals it is the primary means of sensing the environment. The chemoreceptors underlying this process have a rich evolutionary history which mirrors the specialization for diverse habitats across species [1–5]. These proteins evolve with dizzying speed, evincing their position at the interface between the organism and an ever-changing chemical landscape [6, 7]. Surprisingly, however, the olfactory circuit in which these receptors are embedded has deep similarities across vertebrates and invertebrates [8, 9]. In both lineages, organisms have evolved a large repertoire of receptors, many of them broadly tuned (though some receptors are specialists for a given ligand of high importance) [10, 11]. Each primary sensory neuron typically expresses just one receptor, and only neurons expressing the same receptor converge onto a given olfactory glomerulus.

The transcriptional [12, 13] and wiring [14, 15] mechanisms for achieving this circuit organization vary widely across animals. This diversity suggests that these three motifs (broadly-tuned receptors, one neuron-one receptor, and glomerular convergence) are the result of strong selective pressure, rather than evolutionary chance or molecular constraints. Why might this architecture be optimal?

To answer this question, we must specify an objective. Organisms rely on olfaction for many of their basic needs, including mating, feeding, and avoiding predation. From a computational perspective, this means that animals need to solve a wide array of tasks, including discriminating odor identity [11, 16, 17], segmenting odor landscapes [18–22], and matching single odorants to a known mixture (“pattern completion”) [23]. Complicating the picture further are the rich temporal dynamics of olfactory stimuli, which induce correspondingly rich dynamics even in the earliest levels of sensory processing [17, 18, 24–27].

In light of this complexity, we consider a task-agnostic objective: maximizing mutual information. The idea that early sensory processing is organized to maximize mutual information between stimulus and neural representation is known as efficient coding [28, 29]. Of course it is true that in practice, animals benefit from discarding irrelevant information. But critically, the relevance of information is often context- and task-dependent. Since higher areas of the animal brain can only access the olfactory stimulus through the glomeruli, it seems likely that up to the glomerular layer, information maximization is a plausible approximate objective. Many previous works have applied efficient coding principles to olfaction under this reasoning [30–35].

In this paper, we build on previous models for olfactory stimuli and receptor activity [30, 32] to capture the relevant biophysics (such as receptor saturation and widely-distributed odorant concentrations) of the circuit. To allow tractable analysis, we ignore temporal response dynamics in this work (but see the Discussion). Within this framework, we leverage recent technical advances from the domain of unsupervised representation learning to numerically maximize a variational lower bound for the mutual information between the stimulus and the corresponding neural response. We do this with respect to three matrices: the receptor by odorant sensing matrix *W*, the neuron by receptor expression matrix *E*, and the glomerulus by neuron connectivity matrix *G* (see Figure 1). In order to better respect the different timescales at play in the evolution and development of the circuit, we optimize one layer at a time, though joint optimizations yielded similar results. We find that efficient coding recovers the motifs of olfactory architecture that have now been described in several species separated by hundreds of millions of years of evolutionary time [9, 12, 13, 36, 37]. Thus, this striking example of convergent evolution can be understood through the lens of optimality for information processing.

**FIG. 1.**
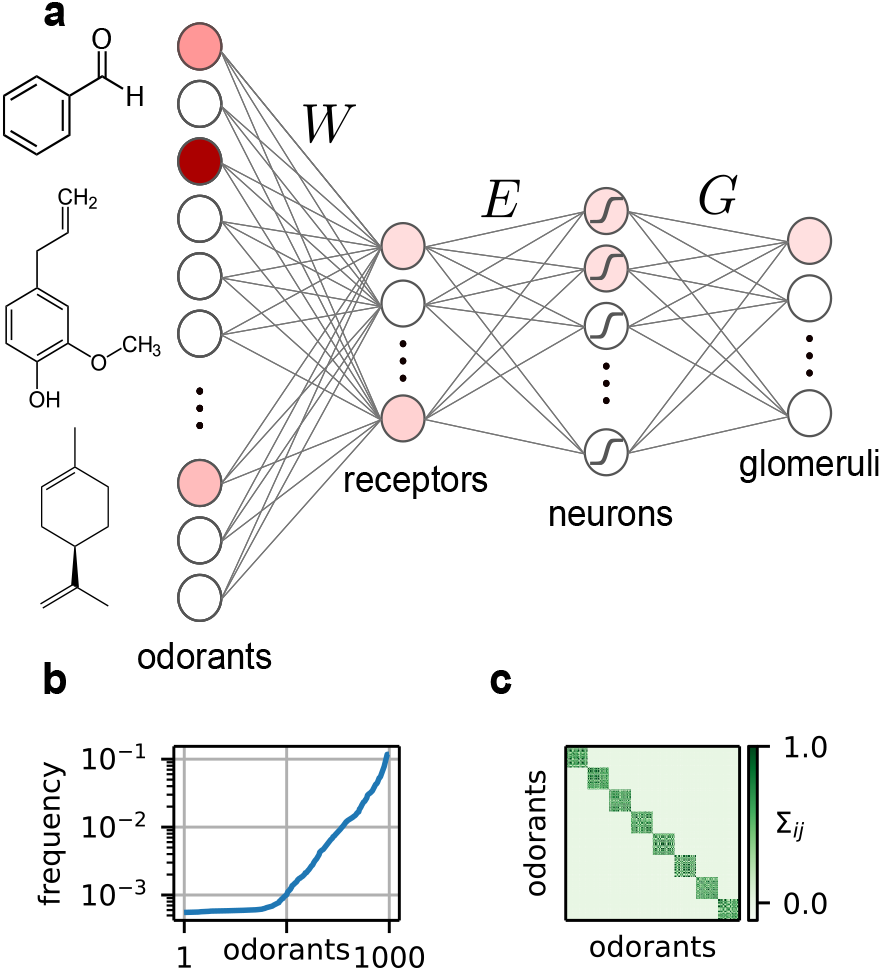
Overview of the model. (a) The structure of the stimulus and the network. (b) The frequency of each odorant across samples. (c) An example covariance matrix of the binarized odorants. The number of blocks (each representing a source) is a parameter we vary—see Section III B.

## II. SETUP

### A. Stimulus and encoding model

We model the olfactory stimulus *c* ∈ ℝ^*N*^ as a sparse vector in the space of concentrations of *N* monomolecular odorants (Figure 1a). We first generate a binary vector *c*_*bin*_ ∈ {0, 1 }^*N*^ using a Gaussian copula. This allows us to tune the mean and covariance of *c*_*bin*_. Means are drawn from a Gamma distribution, so that some odorants are frequent, but most are rare (Figure 1b). We set the covariance of *c*_*bin*_ to be a block matrix, where each of the *k* blocks represents a mixture emitting correlated odorants (Figure 1c). A more realistic covariance matrix would have a nested, hierarchical structure rather than merely a single level of blocks [38], but it is less obvious how to parametrize such matrices—see the Discussion.

We then assign an independent and identically distributed log-normal concentration with variance 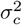 to each odorant in the sample to generate *c* ∈ ℝ^*N*^. A version of this sparse, log-normal model for *c* was used by Qin *et al*. [30], and we extend their model by incorporating varying frequencies, structured covariance, and a degree of sparsity that changes across samples. The circuit encounters one *c* vector at a time, and its task is to compute a useful representation of *c* over the distribution of possible stimuli *p*(*c*). We include full details of the statistical model in the Methods.

Note that while we preserve the presence/absence statistics across samples (so that odorant *α* can occur more frequently than odorant *β*), we do not preserve odorant concentration ratios across samples (odorant *α* cannot occur at a typically higher concentration than odorant *β*). Such differences in concentration can be a critical aspect of identity coding [39]. However, this structure is liable to disruption by the turbulent transport of volatile molecules to the sensory epithelium, as well as differences in molecular diffusivity [39, 40]. Neglecting this structure greatly reduces the number of tunable parameters in our stimulus model.

Given the stimulus, the encoding model is

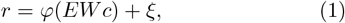

where *r* ∈ ℝ^*L*^ is the firing rate of the neurons,

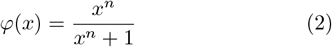

is a Hill function with coefficient *n*, applied element-wise, 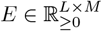 is the neuron by receptor expression matrix, 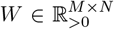 is the receptor by odorant affinity matrix, and *ξ* ∼ 𝒩 (0, *σ*_0_). This Hill function nonlinearity is consistent with measurements in the fly larva by Si *et al*. [41], in which the authors found that a Hill coefficient of *n* = 1.46 best fit their data. The coefficient can be higher in vertebrates (on average *n ≈* 2), where olfactory receptors are G protein-coupled receptors rather than ligand-gated ion channels [42, 43] (but see Grosmaitre *et al*. [44] for *n ≈* 1 in vitro). We confirmed that the qualitative features of the optimal *W* and the optimal *E* did not depend on this coefficient (*SI Appendix: Fig. S1*). Note that, while receptor affinities can be negative [42, 45, 46], we have chosen not to model this inhibition in this work. We discuss this and other limitations of our model, as well as possible improvements, in the Discussion section. While both *W* [30, 32] and *E* [31] have been studied using efficient coding, most works have not considered the interplay between the two (with the exception of Lienkaemper *et al*. [47]).

Finally, the neural activity *r* ∈ ℝ^*L*^ is integrated in regions of neuropil known as glomeruli *g* ∈ ℝ^*M*^, with *M* ≪ *L*. For simplicity, we match the number of glomeruli to the number of receptors, although in reality there could be more glomeruli (as in mice [48]) or fewer (as in mosquitoes [49]). Here the vector *g* consists of one representative dendrite per glomerulus. In paired recordings from pre- and post-synaptic neurons at the glomerulus in *Drosophila melanogaster*, Bhandawat *et al*. [50] found that the transfer function was sigmoid in shape, so we parametrize it using the hyperbolic tangent (again applied point-wise) with varying gain *α*:

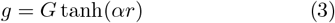

Here, *G* is a connectivity matrix describing the projections between neurons and glomeruli.

### B. Layer-wise efficient coding

Given this model, we perform three separate optimizations. The first is over receptor sensitivities:

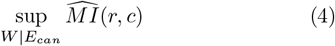

The notation sup indicates that we are finding the matrix *W* that maximizes 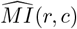. When optimizing *W*, we constrain its elements to be positive. By *E*_*can*_, we indicate that we are plugging in a canonical (one neuron-one receptor) *E* matrix into (1) and holding it fixed to optimize over *W*. Thus we are modeling the ongoing evolution of receptors in the context of the widespread one neuron-one receptor motif, rather than the joint emergence of the two phenomena. While we will largely focus on this layer-wise approach, which allows us to flexibly test null models for each parameter matrix, we also tested joint optimization over *W* and *E*. This joint optimization yielded very similar results to the separate optimizations in (4) and (5) (*SI Appendix: Fig S2*). See Methods for details of the optimization procedures for all three layers).

The next optimization we perform is

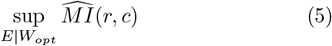

in which we first optimize *W* as in (4), then optimize over *E*. We also shuffle *W* to ensure that our conclusions are robust to the details of the receptor matrix. Varying the details of *W* and the parameters of the environment allows us to probe why canonical expression is so common, as well as exploring the phenomenon of noncanonical expression, as recently characterized in the *Aedes aegypti* mosquito [49, 51]. We constrain *E* to be positive and to sum row-wise to unity, since neurons must allocate a finite budget of receptor expression [49, 51–54].

Finally, we optimize the glomerular layer:

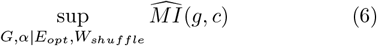

where *W*_*shuffle*_ is a shuffled version of *W*_*opt*_. This is likely to be a more realistic model of true receptors than *W*_*opt*_ due to the biophysical constraints of tuning in real receptors (something we discuss at length in Section III III B). Again we constrain *G* to be positive and to sum row-wise to unity, since glomeruli receive excitatory inputs from finitely many olfactory receptor neurons [8].

We consider the trade-offs of this layer-wise approach, its alternatives, and its biological interpretation in the Discussion.

### C. Maximizing mutual information by proxy

The primary challenge in any efficient coding analysis is the estimation of mutual information. To enable analytical progress, strong assumptions on the stimulus and encoding are required [30, 31, 47, 55–57]. Most simply, if one assumes linear processing of a Gaussian stimulus, the mutual information can be computed in terms of the log determinants of the resulting covariance matrices. However, it is challenging to analytically compute mutual information in more realistic non-Gaussian models like the one we adopt here.

Numerical estimation of mutual information in high dimensions is also fraught with difficulties [56, 58]. Here, inspired by work on deep representation learning, we adopt a conservative approach based on the principle that finding an optimal encoding model does not require an estimate of the precise value of the mutual information. Instead, it is sufficient to maximize a reliably-estimable proxy objective function [59, 60]. Committing to this approach limits our ability to compare different optimized models, but it allows for a much more robust and efficient optimization procedure.

As detailed in the Methods, we leverage a variational formulation for mutual information maximization first proposed by Nowozin *et al*. [61]. Specifically, we maximize a bound on the Jensen-Shannon divergence (JSD) between the joint and marginal distributions of the stimulus and the encoding. This was inspired by previous works showing that the JSD can be estimated more reliably than the Kullback–Leibler divergence (KLD) that defines the mutual information [59, 60]. We describe the trade-offs of this approach in the Methods. We handle constraints on *W, E*, and *G* using the framework of mirror descent; again, see the Methods for details.

## III. RESULTS

### A. Receptors

We first optimized the mutual information over the sensing matrix *W*. Each row of this matrix represents a receptor, and each column an odorant. Thus *W*_*ij*_ is the sensitivity of the *i*-th receptor to the *j*-th odorant. It is helpful to consider that chemoreceptors are both ancient (predating neurons, for example), and fast to evolve— indeed, they are some of the fastest evolving proteins in many organisms [2, 6, 7, 62]. This may be because their conserved function is less critical for survival than that of metabolic or structural proteins. Furthermore, there is very direct evolutionary pressure to modify chemoreceptors in response to changes in the organism’s chemical environment.

Thus it seems that the matrix *W* is relatively tunable (notwithstanding biophysical constraints—see Discussion), and so we asked: to what extent can efficient coding recover the qualitative features of receptor affinities measured in biological circuits? To guide our analysis, we turned to published data on the well-characterized olfactory system of the *Drosophila melanogaster* larva, which has just 21 olfactory receptor neurons, each identified by the expression of a unique receptor. The tractable size of this system permits comprehensive interrogation of the complete receptor repertoire, which was conducted by Si *et al*. [41]. Plugging in a canonical expression pattern *E* as in equation (4), we optimized mutual information between neural activity *r* and the stimulus *c*, using parameters that matched the fly larva circuit.

The resulting *W*_*opt*_ had a non-negligible degree of sparsity (≈ 40% entries functionally 0; see discussion in *SI Appendix A*), and its non-zero elements were well-fit by a log-normal distribution—see *SI Appendix: Figs. S3 and S4*. Both of these features have been derived analytically in previous theoretical work [30, 32] in the limit of vanishing neural noise. Singh *et al*. [34] further argue how sparsity and broad tuning can emerge from simple theoretical considerations about the dimensionality of odor and receptor space. Here we confirmed that these results still approximately hold when neural noise is non-negligible. We next sought to compare our optimized *W* to experimental results in a more detailed and biology-focused analysis, as shown in Figure 2.

**FIG. 2.**
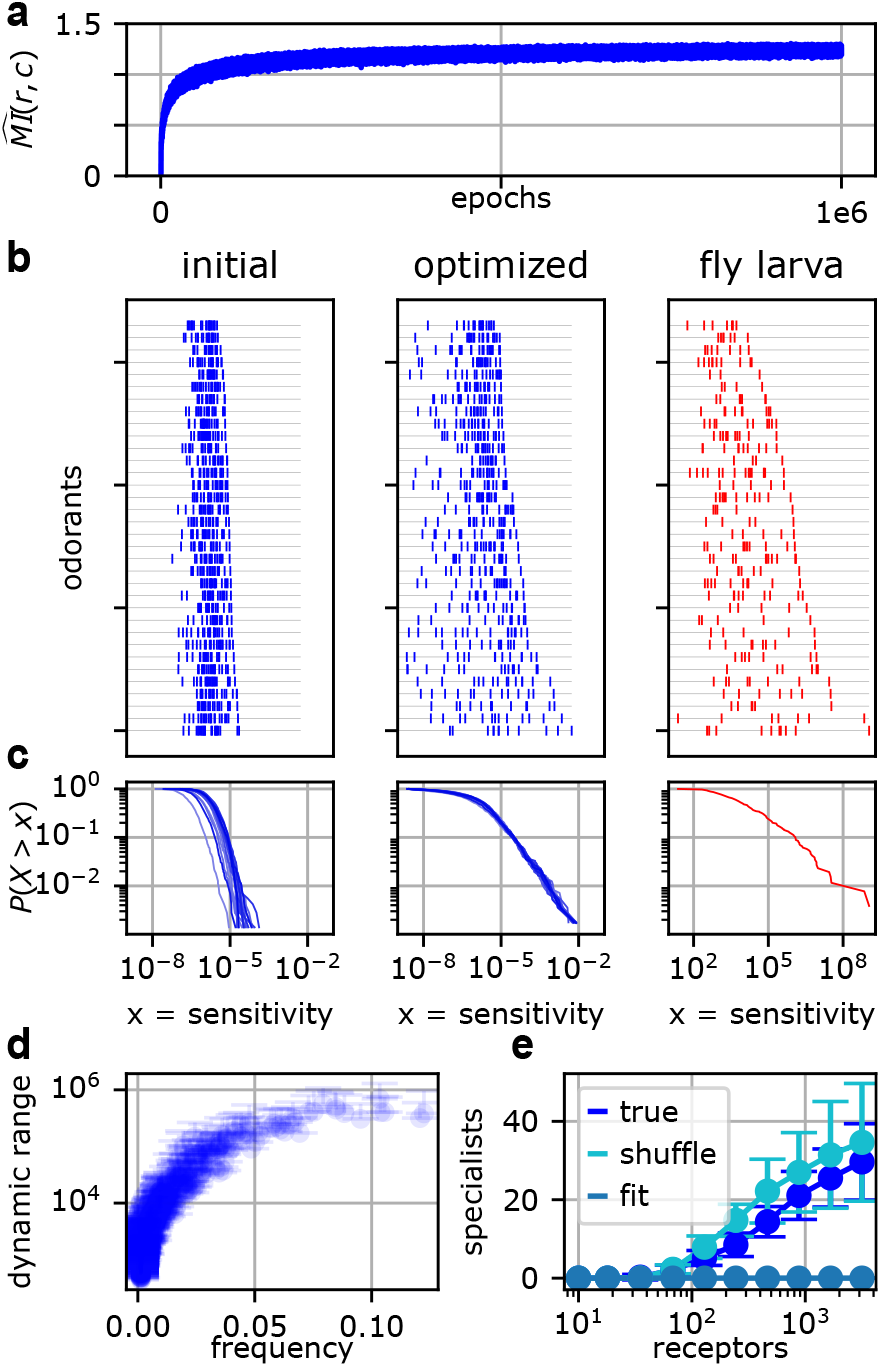
Optimizing over *W* for parameters typical of the fly larva, given canonical expression. (a) The trajectory of mutual information over the course of the optimization. (b) Sensitivity per odorant in the optimization (blue) and fly larva (red). Each tick denotes the sensitivity of one receptor for that odorant. In both cases, there are typically many receptors per odorant, and their sensitivities span several orders of magnitude. Odorants are sorted by their maximum sensitivity within each panel. (c) Distribution of *W*_*ij*_. For the optimizations, 20 runs with different random seeds are shown. More frequent odorants are afforded greater dynamic range. Error bars denote standard deviation over 20 runs (only the upper error bars are shown). (e) For fixed *N* = 1000, the number of “specialist receptors” grows with the number of receptors. “Fit” is an analytic control obtained by fitting a log normal distribution to the optimized W matrix. Experimental data in panels (b) and (c) is from Si *et al*. [41]. Environmental parameters were set to *σ*_*c*_ = 3 and 32 sources.

We found three qualitative matches between our optimized *W* and the fly larva *W*. First, the distribution of sensitivities across all receptors and odorants is broad, spanning approximately six orders of magnitude (see Figure 2c). This heavy-tailed distribution is the object of much analysis by Si *et al*. [41], who argue for the computational advantages of such a code. In our results, it is difficult to confirm a precise power law to the exclusion of other heavy-tailed distributions. But the qualitative agreement shown in Figure 2c may further support the optimality of this distribution for the entries of *W*.

Second, each odorant typically has at least several receptors tuned to it, as shown in Figure 2b. This is necessary to transmit faithful information about the stimulus, since the two orders of magnitude spanned by each receptor’s Hill function activity do not cover the full range of odorant concentrations. (For Figure 2, odorant concentrations were drawn from a log-normal distribution with *σ*_*c*_ = 3.)

Third, some odorants are sensed by a greater number and range of receptors than others (compare the top and bottom lines of Figure 2b). Within the context of our optimization, there is an intuitive reason for this: it is the more frequent odorants that are afforded greater dynamic range, as shown in Figure 2d. While this result may hold in experimentally measured receptor sensitivities, testing it directly would require quantitative knowledge about natural olfactory landscapes, which are notoriously difficult to characterize.

More fundamentally, however, this last finding underscores one limitation of our efficient coding framework. Mutual information is a statistical measure of the dependence between two distributions and does not privilege any particular component of the stimulus for reasons of ecology or behavior. Thus, when optimized, our model circuit simply prioritizes sensing of common odorants (although it is important to note that this is not a generic prediction of efficient coding [31, 63, 64]). In natural settings, by contrast, animals may have evolved receptors to sense an infrequent but critically important odorant (such as a toxin or pheromone) with high sensitivity across a wide range of concentrations, as in Sakurai *et al*. [65]. Thus, although the trend in Figure 2d may hold on average, it is likely to admit important exceptions in biological circuits.

The latter observations consider the circuit’s ability to report information about a given odorant (studying columns of *W*). The complementary viewpoint is to begin with a receptor and study its tuning across odorants (rows of *W*). For example, much of the recent experimental work on olfactory receptors has emphasized the promiscuity of their binding. Rather than a “lock and key” pairing, ligands were found to fit loosely in the binding pocket of the insect receptor *Mh*OR5 [66]. This strategy is partly due to the odorant-receptor bottleneck (*N* ≫ *M*), and under our model, most receptors indeed exhibit broad tuning (*SI Appendix: Fig. S3*). However, this bottleneck argument does not explain why some receptors seem to have narrow tuning for one or a handful of odorants [10], especially in organisms with greater numbers of receptors (such as in mice, where *M* ≈ 1300).

To explore this question, we varied the degree of the bottleneck (keeping *N* fixed and increasing *M*) and inspected the resulting optimized receptor profiles. As we increased the number of receptors, the circuit acquired an increasing number of “specialist receptors” (see Figure 2e). We formalized this using the following criterion: if the maximum sensitivity of a receptor for an odorant was at least two orders of magnitude greater than the 99th percentile sensitivity, then we counted it as a specialist for that odorant. This corresponds to testing a panel of 100 odorants, and finding that a receptor has 100x greater sensitivity for its specialized odorant than for all other odorants in the panel.

In Figure 2e, we plot two important controls. One is a shuffled *W*. For this control, the trend still holds, which indicates that specialist receptors are developed by pushing up the sensitivity of the specialist receptor-ligand pair, rather than pushing down the sensitivities of the specialist receptor for other ligands. Therefore, a specialist in our model is a receptor with mostly typical sensitivities, and one very high outlier.

Our other control is an analytic fit. Here we fit a log-normal distribution to the non-zero values *W*_*ij*_ for each value of *M*, then sample from that distribution to generate a *W*, and count specialists in that *W*. While the log-normal is heavy-tailed and fits *p*(*W*_*ij*_) well for the bulk of the distribution (*SI Appendix: Fig. S4*), it does not generate the extreme outliers needed to produce any specialist receptors under our very stringent criterion. This demonstrates that the effect is not driven simply by drawing more samples from the same distribution.

These findings suggest that specialist receptors may “look normal” until their target ligand is found. Conversely, given knowledge about a specialist receptor, it may be worth testing other odorants against that receptor to see if they are detected at reasonable concentrations (such as in Meyerhof *et al*. [67]).

Interestingly, despite the allocation of greater dynamic range to higher frequency odorants (Figure 2d), we did not find that they were more likely to be targeted by specialized receptors (*SI Appendix: Fig. S5*). This may reflect the fact that, when specialist receptors are saturated in the presence of their target odorant, they effectively exacerbate the bottleneck faced by the rest of the circuit 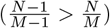. This suggests that there may be slight pressure against developing specialist receptors for common odorants.

In this way, optimal sensing matrices *W* may leverage structure in the stimulus, as well as promiscuous binding, to get useful information from a specialist receptor when the target odorant is not present. Some support for this idea can be found in studies [68, 69] which showed that background odors can compromise the detection of a ligand by its cognate specialist receptor. Conversely, host plant volatiles were shown to synergistically improve pheromone detection in the silkmoth *Bombyx mori* at the receptor neuron level [70].

Finally, one important check on our numerical procedure was to confirm that the distribution of the optimal *W* does not depend on initialization. In Figure 2, the initial *W* is a scaled log-normal: *W*_*init*_ = 1*/*𝔼 [ ||*c*||]*e*^*Z*^, where 𝔼 [||*c*||] is the mean magnitude of the stimulus vector (see Methods). We confirmed that scaling the log-normal to the minimum sensitivity instead, below which *W*_*ij*_ is effectively 0, did not change the results in Figure 2. (We derive this value and discuss its implications in *SI Appendix A*.)

Initialization had virtually no effect on the distribution of *W*_*opt*_ (as in panels b,c) or the relationship between frequency and dynamic range (panel d). The one instance of dependence on initialization was seen in the high *M* regime (*M >* 500) in panel e. With so many receptors, the bulk of *W*_*ij*_ could remain close to initialization without degrading performance. This increased the number of specialists that emerged by roughly a factor of 2 (maximum 57 rather than 27 as in Figure 2), but did not change the trend. We plot Figure 2 for this alternative initialization in the *SI Appendix: Fig. S6*.

To summarize, the *W*_*opt*_ which results from our optimization procedure shares a number of properties with the experimental *W* measured in the fly larva. The distribution of values *p*(*W*_*ij*_) spans six orders of magnitude and the non-zero values are well fit by a log-normal density. Receptor tuning is surprisingly broad, given that we do not explicitly model the imperfect binding of real ligand-receptor pairs. Lastly, increasing *M* permits the emergence of specialists in circuits with larger receptor families.

### B. Expression

Perhaps the most striking feature of canonical olfaction is the “one neuron-one receptor” rule. From a computational perspective, it is not clear why this should be optimal, and yet it has emerged in various organisms including flies [37], mice [12], and ants [13]. Notably, each of these animals has developed an entirely different mechanism to generate this pattern of expression.

In flies, a core set of 15-20 transcription factors and cisregulatory elements acts combinatorially to establish the expression of a single odorant receptor (OR) in a given cell type [71]. In mice, on the other hand, the OR loci from 18 chromosomes come together in a hub that chooses a single receptor allele for expression and silences all the rest [12]. Very recent work has demonstrated how a complex gene expression program whose usage varies tightly with position in the olfactory epithelium orchestrates this OR choice [53, 54]. In ants, many odorant receptors are transcribed, but only the most upstream gene is exported out of the nucleus [13, 72].

These and other disparate mechanisms [71] suggest that the one neuron-one receptor rule may confer a strong fitness advantage. Accordingly, when we numerically maximized our proxy mutual information with respect to *E* in (1) for numbers typical of the fly larva, we found that a clear pattern of canonical expression emerged, as shown in Figure 3. This was despite random initialization (Figure 3b).

**FIG. 3.**
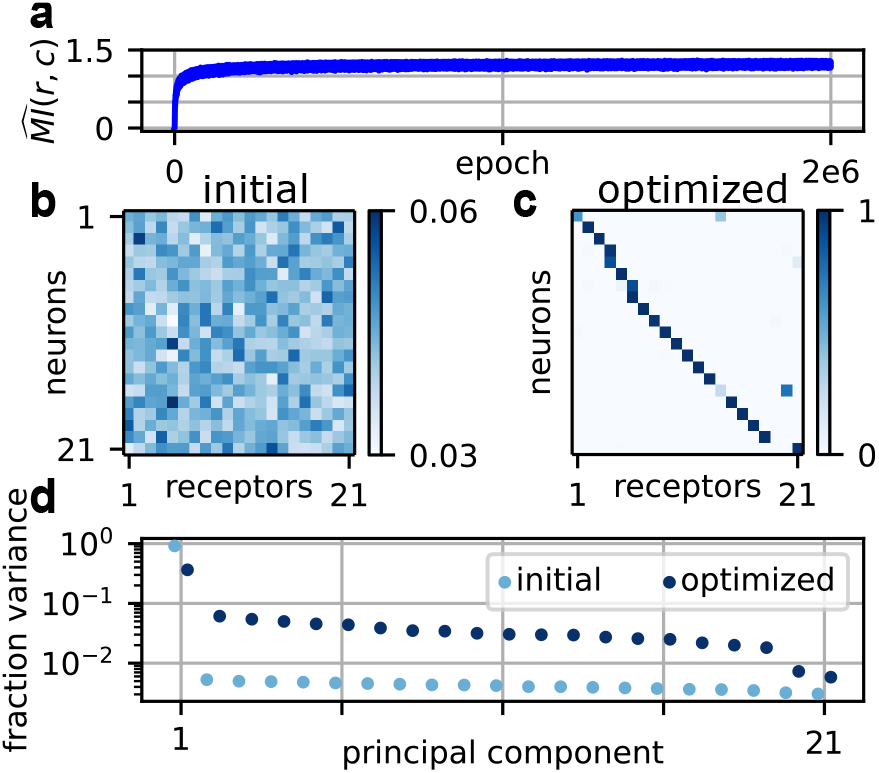
Optimization over *E* for numbers typical of the fly larva. (a) The trajectory of mutual information over the course of the optimization. (b) The initial expression matrix. (c) The optimized expression matrix. The optimization is over *E* in (1), plugging in the optimal *W* from Figure 2. (d) The fraction of variance in neural activity *r* explained by each of the top principal components, given the initial expression from panel (b) and the optimized expression from panel (c). Environmental parameters were set to *σ*_*c*_ = 3 and 32 sources. Statistics are computed over 1000 samples.

Immediately, however, we are confronted with a “chicken-or-egg” problem. In (4), we optimize *W* given canonical *E*, then in (5) we optimize *E* with the resulting *W*_*opt*_. This could in principle bias the optimization over *E* to converge on canonical expression. To account for this possibility, in our subsequent analysis we optimize *E* given five plausible alternative models for *W* (see analysis below, and *SI Appendix: Fig. S13*, for details). We find that each of these models favors canonical expression, which indicates that the result is not merely an artifact of our layer-wise approach.

Given that the solution is robust, why is canonical expression optimal? In the limit of low neural noise, one simple way to increase mutual information is simply to decorrelate activity, when a distribution of activity is computed over the stimulus distribution *p*(*c*). This is a standard prediction of many efficient coding analyses [29, 73]. Accordingly, we find that the distribution of variances explained by the principal components of activity is flatter under canonical expression than under random expression (Figure 3d). The best way to decorrelate activity in this way is to place neurons at the corners of the simplex in gene expression space, since total receptor expression per neuron must sum to unity. This corresponds precisely to the one neuron-one receptor rule. (See Lienkaemper *et al*. [47] for a related analysis in the case of one neuron.)

We next sought to understand if the same result would hold in a larger system. We chose a scale comparable to the adult fly, with *M* = 60 receptors and *L* = 1260 olfactory neurons. This is a natural focus, since olfactory processing in the fly is well characterized (see for example Hallem and Carlson [11]), and these numbers permit reasonable numerics. We first optimized *W*, and found a qualitative match to the measurements in Hallem and Carlson [11], with most receptors exhibiting broad tuning (*SI Appendix: Fig. S7*).

Surprisingly, when we optimized over *E* using this *W* matrix, a handful of receptors were typically coexpressed per neuron as shown in Figure 4a. Note that here, and throughout this paper unless otherwise noted, we have initialized expression to be noncanonical, in order to test the robustness of canonical expression. When we initialize expression to be canonical instead, we obtain canonical expression as a solution (see *SI Appendix: Fig S9*). As a result, we can conclude that both canonical and non-canonical expression are local solutions to the information-maximization problem in this setting.

**FIG. 4.**
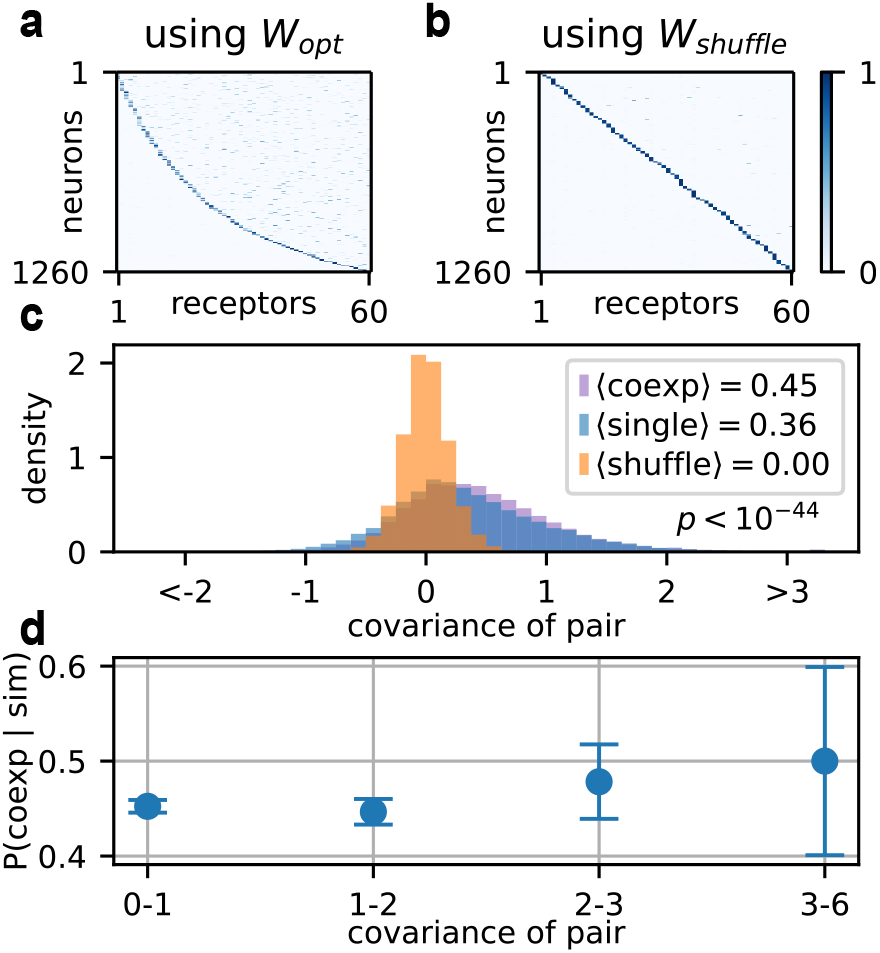
Optimization over *E* for numbers typical of adult *Drosophila*. (a) The optimized expression *E* obtained from plugging in the optimized sensing matrix *W*_*opt*_. (b) The optimized *E* obtained from plugging in a shuffled version of *W*_*opt*_. (c) Covariance between *W*_*opt*_ receptor pairs which are coexpressed (purple), *W*_*opt*_ singly expressed pairs (blue), and *W*_*shuffle*_ receptor pairs (orange). Covariances are computed on log sensitivities. (d) Probability of coexpression given binned covariances. For panels (a) and (b), environmental parameters were set to *σ*_*c*_ = 2 and 64 sources. Data in (c) and (d) are across 21 runs with varying environmental parameters *σ*_*c*_ and Σ_*c*_, given by the bottom three rows of the phase diagram in Figure 5. The p-value for the difference in distribution between coexpressed pairs (purple, *n*_*c*_ = 16038) and singly expressed pairs (blue, *n*_*s*_ = 21132) is obtained using the Mann-Whitney U test. The error bars in panel d are computed using the Wilson score interval with *α* = 95%.

Interestingly, recent work in the *Aedes aegypti* mosquito has discovered receptor coexpression in many olfactory receptor neurons (ORNs) [49, 51]. Reanalysis of existing data by Adavi *et al*. [51] further characterized occasional exceptions to the one neuron-one receptor rule in *Drosophila*. The extent of this coexpression is controversial [71, 74] but some unambiguous cases have been known for many years [75, 76].

It is important to note that, in both *Aedes aegypti* and *Drosophila*, the majority of coexpressed chemoreceptors are disproportionately close to each other in genomic and phylogenetic space [51]. Adavi *et al*. [51] term this “coexpression by descent.” In only a minority of cases did the authors find that coexpressed receptors were far from each other (“coexpression by co-option.”) This suggests that much of the phenomenon may be a by-product of shared gene regulation. Since families of chemoreceptors frequently expand via tandem duplications (the “birth and death model” of gene evolution) [7, 77], there are many examples of neighboring ORs with similar sequences whose coexpression might be driven by cis-regulatory elements [71].

These molecular mechanisms are unlikely to be fine-tuned for optimal information processing. However, the fact remains that different insects (like *Drosophila* and *Aedes aegypti*) exhibit strikingly different levels of coexpression. To understand the functional significance of these differences, it is helpful to consider the similarity between receptor affinity profiles, as shown in Figure 4c and d. Under this reasoning, coexpression is primarily driven by shared regulatory machinery, but once it emerges, its fitness effect depends on the similarity of the coexpressed receptors, with coexpression of more similar receptors being less detrimental. Since phylogenetically related receptors typically have similar affinity profiles [78, 79], these results are consistent with the analysis of Adavi *et al*. [51].

Given these subtleties, why did we obtain such a clean canonical expression profile in our analysis of the larval fly (Figure 3)? The answer may be in the number of receptors *M*. Sweeping over this parameter, we found that optimized receptor matrices with small *M* were much closer to full rank (*SI Appendix: Fig. S8*), indicating that their receptors were more differentiated. These results suggest that organisms with only a handful of evolutionarily mature receptors cannot afford similar affinity profiles.

Our findings changed qualitatively between the scales corresponding to the fly larva (Figure 3, *M* = 21) and to the adult fly (Figure 4, *M* = 60). To better understand this puzzling result, we inspected the optimized matrix *W*_*opt*_ in the adult fly setting. We found a distinctive low rank structure in *W*_*opt*_ across environmental parameters that was especially pronounced for higher values of *σ*_*c*_ (*SI Appendix: Fig. S10*). This is driven by our frequency model since, as indicated in Figure 1b, only a small subset of odorants occur with high frequencies, while the bulk occur at low frequencies. Flattening the frequency distribution abolished the low rank structure (*SI Appendix: Fig. S11*). This degeneracy in *W*_*opt*_ permits either canonical or noncanonical olfaction, suggesting that the precise details of expression do not matter when *W* is low rank.

While potentially informative, this result also reflects a limitation of our mutual information approach. Under this objective, high-frequency odorants are allocated greater dynamic range (Fig. 2d), which induces a lowrank structure in *W*_*opt*_. In reality, organisms may need to detect common and rare odorants with equal sensitivity. Additionally, biophysical constraints limit the number of ligands a receptor can bind. Indeed, Si *et al*. [41] found that covariance in olfactory sensory neuron activity is primarily driven by the geometric structure of odorants. This geometry constitutes a significant constraint on receptor affinities that is not accounted for in our model, and thus real receptors are very likely to be less finely tuned than our *W*_*opt*_ matrices.

In light of these limitations, we next studied expression using a shuffled version *W*_*shuffle*_ of the optimized *W* matrix, as in Figure 4b. Shuffling allowed us to preserve the scale, distribution, and sparsity of *W*_*opt*_, while breaking the low rank structure. It also accounts, albeit coarsely, for the suboptimal tuning of real receptors. Plugging in *W*_*shuffle*_ favored more canonical expression, as seen in Figure 4b.

We next sought to understand when the one neuron-one receptor rule emerges at a scale closer to the number of receptor types found in mammals such as humans (*M* = 400) and mice (*M* = 1300). Optimizing *W* and *E* for *M* = 400, we found very similar results compared with the *M* = 60 case; for *W*_*opt*_, coexpression is tolerated, but for *W*_*shuffle*_, near-perfect single expression emerges (see *SI Appendix: Fig. S12*; we did not optimize over *M* = 1300 due to memory constraints). Critically, therefore, our results indicate that a larger number of receptors does not drive coexpression. Instead, it is the extremely low-rank structure of the optimized *W* matrix that permits (but does not favor) coexpression.

Real *W* matrices are most likely best modeled by something between *W*_*opt*_ and *W*_*shuffle*_. They have some correlation structure, due to the development of new odorant receptors from existing ones, but are unlikely to be extremely low rank. We therefore implemented two more models for *W* : a log normal analytic *W* with block covariance structure, to model families of related receptors, and a log normal *W* with Toeplitz covariance structure, to model a range of similarities across receptors in the adult fly. We initialized expression to simulate coexpression by descent; that is, coexpression of correlated receptors. However, we found that canonical expression was still largely favored regardless of receptor details (see *SI Appendix: Fig. S13*).

To ensure that these results did not depend on specific parameter values, we next swept over the two important parameters of our odorant model: the structure of the covariance matrix Σ_*c*_ and the log-normal noise parameter *σ*_*c*_. We parameterized Σ_*c*_ by giving it a block structure and varying the number of blocks (see Figure 1 and the Methods). Ecologically, each block of correlated odorants corresponds to a source in the organism’s environment. Precise quantification of the natural olfactory environment is elusive (see Yang *et al*. [80] and Zhou *et al*. [38] for recent efforts in the fly context), but it is plausibly contained in the large space we explore.

Sweeping across these parameters did not reveal any strong dependence of our results on environmental statistics (Figure 5). For *W*_*opt*_, we did not observe significant changes in *E*, irrespective of whether we initialized with a non-canonical expression pattern (Figure 5a) or canonical expression (*SI Appendix: Fig. S9*). We can therefore conclude that with the low-rank *W*_*opt*_ neither solution is strongly favored, regardless of environmental structure. For *W*_*shuffle*_, however, we found that near-canonical expression emerged across almost all combinations of environmental parameters (as shown in Figure 5b), despite noncanonical initialization. We found only a weak dependence on environmental parameters given *W*_*shuffle*_, and the trends were reversed compared to *W*_*opt*_. We tested two other unstructured models for W: shuffling *W*_*opt*_ within rows (in order to preserve mean receptor tuning), and fitting a log normal distribution to *W*_*opt*_. Both of these favored canonical expression when *E* was then optimized (see *SI Appendix: Fig. S13*).

**FIG. 5.**
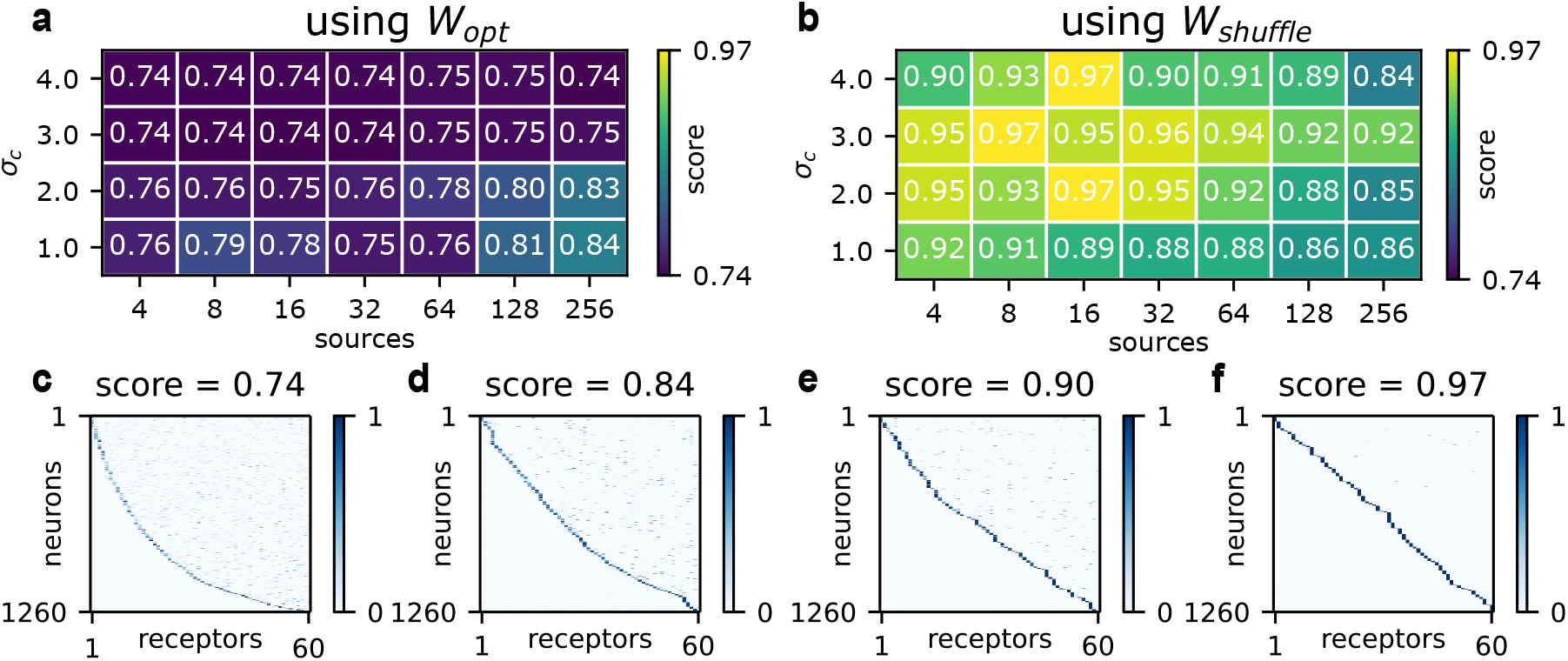
The degree of canonical expression in optimized *E* matrices while varying environmental parameters *σ*_*c*_ and Σ_*c*_. (a) The phase diagram using optimized *W*. (b) The phase diagram using shuffled *W*. (c) through (f): example expression matrices taken from the above phase diagrams. The score is 1 − 𝔼 [*H*(*E*_*i*_)], where *H*(*E*_*i*_) measures the entropy of the *i*-th row of *E*, when the expression is viewed as a probability distribution over receptors, and the expectation is taken over the rows of *E*. Here, we initialize the expression matrices to be noncanonical: each neuron expresses 3-7 receptors at roughly equal levels, for a “canonical score” of 0.74. In SI Appendix: Fig. S9, we show the corresponding results for canonical initialization.

Finally, we sought to understand how the level of neural noise *σ*_0_ affected these results. We swept across *σ*_0_ for a representative pair of environmental parameters (blocks = 64, *σ*_*c*_ = 2) (*SI Appendix: Fig. S14*). We found that at extremely low noise levels (*σ*_0_ = 0.01), both solutions were equally favored, but for more realistic levels (*σ*_0_ ∈ [0.1, 1.0]), canonical olfaction was favored as shown above. All of the above results were run for *σ*_0_ = 0.1, a value we chose because it sets the noise to be on the order of the mean activity (see *SI Appendix: Fig. S16* and Glomerular convergence). Increasing *σ*_0_ beyond this point further favored canonical olfaction. This trend is consistent with the analytical theory of Lienkaemper *et al*. [47] (see Discussion).

In sum, our analysis suggests that receptor co-tuning, rather than environmental statistics, is decisive in determining optimal expression. In our model, only extremely low-rank *W* matrices permit significant levels of noncanonical olfaction. Biologically, such a set of receptors would have highly correlated tuning across many odorants. A number of more plausible models for *W* support largely canonical olfaction for realistic levels of neural and environmental noise. This may explain why the one neuron-one receptor rule has emerged in such distantly related organisms, despite the fact that these organisms sense different chemical environments with receptor families that are accordingly divergent. Conversely, our results suggest that coexpression may emerge when receptor affinities exhibit a large amount of redundancy.

### C. Glomerular convergence

Olfactory sensory neurons converge onto regions of neuropil known as glomeruli [81]. In the canonical model, only neurons expressing the same receptor converge onto a given glomerulus [82]. This “glomerular convergence” confers an intuitive advantage: given canonical expression, the signals from each receptor can be averaged across neurons without mixing across receptors. We plugged in *W*_*shuffle*_, the corresponding *E*_*opt*_, and optimized our proxy for mutual information over *G* and *α* as in (6). We confirmed that glomerular convergence is recapitulated in our model across a wide range of environmental parameters (see Figure 6a-c, and *SI Appendix: Fig. S15* for sweep).

**FIG. 6.**
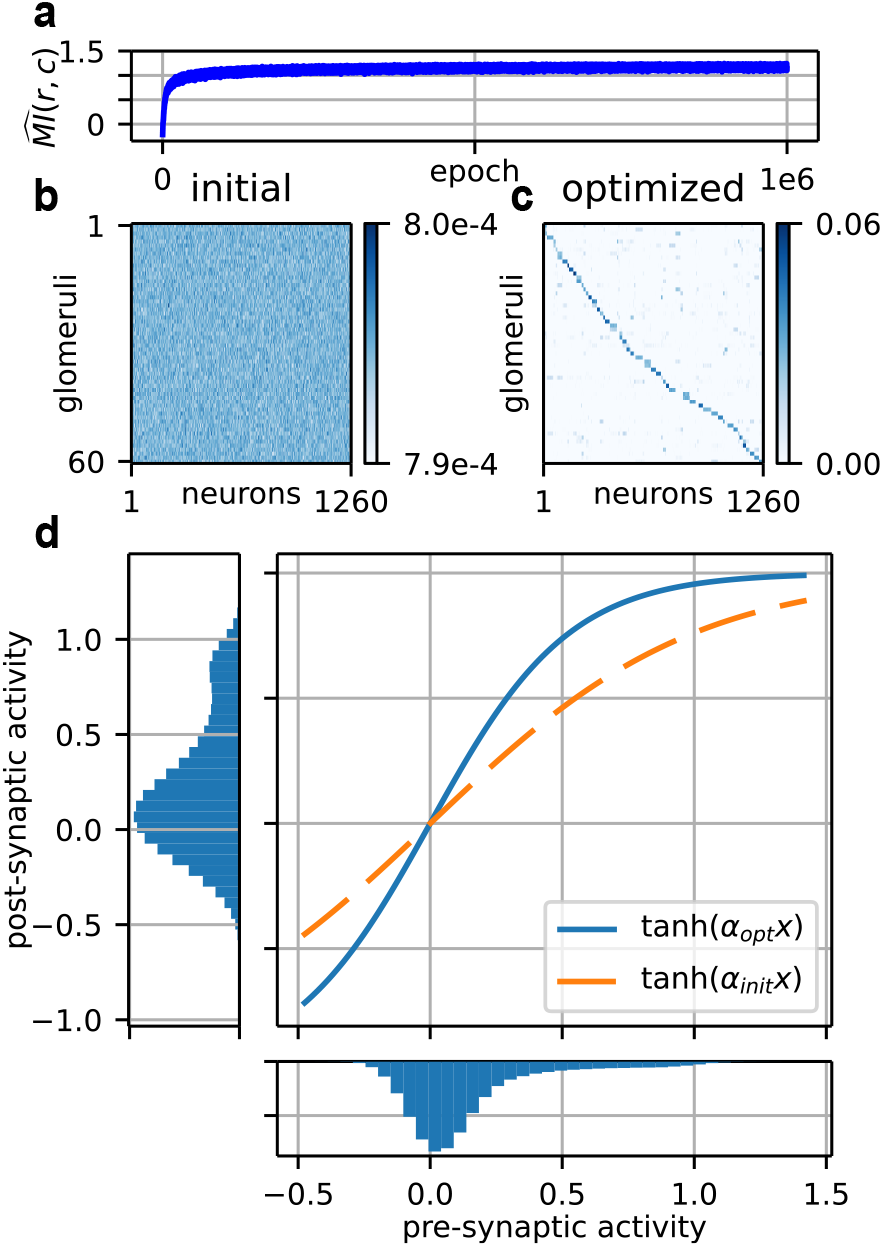
An example optimization over the glomerular layer for 64 sources and *σ*_*c*_ = 3. (a) The course of the mutual information estimate over the optimization. (b) The initial random connectivity. (c) The optimized connectivity. Neurons are sorted by their highest expressed receptor. (d) The transformation at the glomerular layer. A higher *α*_*opt*_ serves to flatten the distribution of post-synaptic activity.

But glomeruli do not just serve to denoise olfactory neuron responses. Careful recording of both the presynaptic ORN and the post-synaptic projection neuron (PN) in *Drosophila* has characterized other aspects of the transformation [50]. Chief among these is “histogram equalization,” in which low ORN activity is amplified but higher activity is not. Since ORN activity is clustered near 0, this induces a more balanced distribution of activity in the post-synaptic neuron, which in turn enhances information transmission. This is a classical prediction from the early days of efficient coding [83].

We first confirmed that our activity clustered near 0 as in Bhandawat *et al*. [50] (see Figure 6d, bottom axis, and *SI Appendix: Fig. S16* for sweep). This is likely necessary due to the log-normally distributed stimulus; if the optimal *W* is to account for the rare, highest concentration odorants, then the bulk of stimuli will generate responses closer to baseline. Accordingly, we found that as we increased *σ*_*c*_, the distribution of activity was more sharply peaked around 0 (*SI Appendix: Fig. S16*).

Optimizing over the gain parameter *α* in (6) resulted in a qualitative match to data from adult *Drosophila*, in which the post-synaptic activity was more evenly distributed across the dynamic range of the neuron (Figure 6d). The resulting histograms were not perfectly equalized, but this may reflect the fact that increasing the entropy of the representation *H*(*g*) addresses just one term out of two in the expression for mutual information:

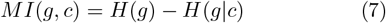

In sum, our results are consistent with the idea that glomerular convergence allows for denoising and histogram equalization of olfactory receptor neuron responses when expression is canonical (although glomerular convergence can also account for variation in the abundance of different cell types–see Discussion). As we previously observed that canonical expression is largely favored across a range of environmental parameters, we did not investigate what connectivity emerges when multiple receptors are expressed in each sensory neuron. However, as long as neural noise is a limiting factor, convergence is the likely solution for denoising.

## IV. DISCUSSION

In this work, we set out to understand why odor coding exhibits deep similarities across vertebrates and invertebrates. Using efficient coding, we recovered three motifs of a widely shared olfactory logic: broad receptors, single receptor expression, and glomerular convergence.

### A. Receptors

The non-zero sparsity of the optimal *W* and the broad distribution of its values have been derived in previous theoretical works that approximate the mutual information *MI*(*r, c*) as the entropy *H*(*r*), an approximation which becomes exact in the limit of no neural noise [30, 32, 34]. Our analysis first recovers these results in the regime where neural noise is non-negligible, then builds on them by studying more granular features of *W*_*opt*_ such as receptor tuning, odorant sensitivity across receptors, and specialists. These details allow us to make new predictions for experiment, particularly regarding specialist receptors.

In our model, receptor tuning is much more promiscuous (most receptors respond to most odorants) than strictly necessary to map all *N* odorants in an approximate labeled-line scheme. This correspondence was surprising, since broadly tuned receptors in biology might simply be a byproduct of ligand-receptor biophysics. If this were the case, then such receptors would not emerge in our unconstrained model when we initialize to zero. Additionally, recent work in the migratory locust *Locusta migratoria* has characterized strikingly narrow receptor tuning [84]. This constitutes compelling evidence that broadly tuned receptor families are not inevitable.

Our results suggest instead that such tuning confers an information processing advantage. The rich theory of compressed sensing may shed light on this advantage, but it is difficult to apply classical results from this field directly to the problem at hand, since a linear measurement model is usually assumed [85]. Recent progress in nonlinear compressed sensing may enable theoretical understanding of the bottleneck problem in olfaction, but much remains unknown [30, 33, 34, 86–88].

These broadly tuned receptors enable a combinatorial coding scheme in which many more odorants can be represented than there are receptors [89]. To the extent that specialists do emerge in our optimizations, they have increased their sensitivity for the target ligand rather than decreasing their sensitivity for off-target ligands, and are thus still available for sensing other odorants.

One prediction of our analysis is therefore that, in the absence of the target ligand, specialist receptors may be co-opted for the sensing of other relevant molecules at realistic concentrations. This idea is indirectly supported by their diminished performance in the presence of background odorants [68, 69], which indicates that specialist receptors preserve some tuning for off-target ligands. Our results suggest that this phenomenon may be a feature, not a bug, when the specialized ligand is not present. This could be further explored simply by presenting specialist receptors with a typical panel of common odorants, ideally ones not naturally co-occuring with the target ligand.

The question of the relevant range of concentrations for odorant stimuli is critical. Wachowiak *et al*. [90] have recently argued that the concentrations used in typical experimental studies may exceed those encountered in the environment by several orders of magnitude. If this is true, than much of the broad tuning of receptors as currently characterized by experiment would be functionally inaccessible to the organism. On the other hand, careful experimental work [91, 92] (see [93] for a review), supported by models of turbulent transport [39, 40], suggests that odorant concentrations fluctuate wildly at the sensory epithelium. Animals may therefore sense odorants quickly during high-concentration bursts, rather than averaging over long timescales. This is supported by behavioral experiments in which flies can execute odor-guided behavior within 100ms [94], as well as the discovery that rodents can identify odors within a similar temporal window [16–18]. Note, however, that these turbulence arguments do not apply to the *Drosophila* larva, since the fly spends most of its larval stage inside a food source [95].

Even granting that experimental concentrations are unnaturally high, the problem is mitigated by the consideration that all studies necessarily use a limited panel of odorants. As more odorants are tested, more extremely high sensitivities will be filled in for each receptor, and the current picture of combinatorial coding might survive, even if all sensitivities are shifted upwards. Our model, which only considers the relative values of sensitivity and concentration, can say little to resolve this question, which amounts to fixing the mean of the stimulus distribution *p*(*c*). But our results do suggest that the variance of *p*(*c*) is likely to be high, since this variance is what drives the spread in the optimized sensitivities, and these in turn are a close match to experimental data.

There are several limitations of our receptor-level analysis. First, we have only accounted for excitation of sensory neurons by odorants. This was primarily because inhibition was not detected by calcium imaging in Si *et al*. [41], which was our primary experimental comparison. In reality, however, antagonistic interactions play a crucial role in mammalian odor coding [42, 45, 46]. Inhibitory responses also emerged in previous theoretical work which optimized over *W* given a non-zero baseline activity in neurons [30]. Mechanistically, a minimal two step model in which the odorant first binds the receptor with some affinity and then activates it with a different, potentially uncoupled affinity, explained a number of classic observations in psychophysical experiments such as synergy and overshadowing between pairs of odorants [96] (see also [97]). In future work, we hope to optimize over the parameters of this more realistic biophysical model and revisit the above analyses (initial attempts to do so proved numerically unstable within our framework).

Second, lurking in any biophysical model is the question of where to place the nonlinearity. In (1), we have placed the expression matrix *E* inside of the Hill function. Therefore we are using the Hill function as an empirical fit to ORN activity in the presence of multiple odorants, not as a mechanistic model for cöoperativity in odorant receptor signaling. A plausible extension of our model would place *E* outside of the nonlinearity, then include another nonlinearity for neural activity. To our knowledge, this issue has not been considered in previous works on efficient coding in olfaction, either because they assumed linear processing of the stimulus [31, 47] or canonical olfaction [30, 32] (i.e., a singly-expressing *E* whose placement therefore has no effect).

Third, on the theoretical side, a significant body of work suggests that early olfaction implements a form of compressed sensing [30, 32, 34, 87, 88]. In this framework, the bulk statistical properties of *W* matter more than the detailed arrangement of particular elements [98]. This was part of our motivation for considering alternative models to the optimized *W*, such as the shuffled and analytic *W* matrices, when analyzing downstream processing. However, biological receptor affinity matrices are subject to odorant-specific evolutionary pressure, as evidenced by the emergence of specialist receptors, and sense different odorants with different dynamic ranges (see Figure 2). These biologically-relevant details can be captured by optimizing *W*, but not by assuming independent random affinities.

Last, and most fundamentally, biological evolution is a much more constrained and local process than our numerical optimization procedure. For example, real receptors likely cannot change their affinity for one odorant without changing their affinities for others, especially since odorant geometry is a primary determinant of receptor specificity (see Si *et al*. [41] and del Mármol *et al*. [66]). Furthermore, organisms develop new receptors from existing ones through a birth-and-death model [7, 77], rather than optimizing a set of randomly initialized receptors as we have done. Any attempt to encode these evolutionary dynamics in our optimization would be computationally challenging, but a principled approach could be fruitful.

With these constraints in mind, it was perhaps surprising that efficient coding could predict any aspects of receptor tuning. Our optimization process becomes more realistic as we compare to more evolutionarily mature receptors that diverged from each other long ago, so the correspondence we found may indicate that the receptors of the larval *Drosophila* are reasonably mature and tunable.

### B. Expression

One key takeaway from our work is that approximate one neuron-one receptor expression emerges across a broad range of environmental parameters and for all but one of the model receptor matrices which we considered. In any normative model, a chief concern is the extent to which such results depend on specific details of the setup. This is why we have swept over a broad range of parameter values for Σ_*c*_, *σ*_*c*_, and *σ*_0_, and inserted multiple models for *W*. The fairly general emergence of canonical expression in these analyses is likely due in part to the decorrelation of activity (see Figure 3d) that occurs when neurons “spread out” in gene expression space by choosing one receptor. A complete answer, however, would require a more developed theory for the nonlinear setting, which we leave for future work.

Critically, we do not find evidence that receptor coexpression should be finely tuned to either the statistics of receptors or the environment. On the contrary, we find that one neuron-one receptor is typically preferred, and that exceptions to this rule should have varying effects depending on receptor similarity. Stronger claims would need to be weighed against the recent work suggesting that many coexpressed pairs are recent duplicates whose coexpression is driven simply by shared regulatory factors [51, 71]. Ramdya and Benton [99] raise the possibility that such instances may reflect a transient evolutionary state, in which recent duplicates are coexpressed only until regulatory machinery has “caught up” to the duplication event by creating new cell types.

In this respect, our findings differ from a very recent theoretical analysis by Lienkaemper *et al*. [47], who characterized the dependence of optimal expression on environmental noise and the “signal correlation” *W* Σ_*c*_*W* ^⊤^ using a linear-Gaussian theory. Discrepancies may be due to the nonlinearity we include in the receptor activity or the log-normality of our odor model. It is also important to note that our formulation differs slightly from theirs: whereas they add noise to a Gaussian stimulus *c* = *c*^′^ + *ξ* and compute *MI*(*r, c*^′^), we compute *MI*(*r, c*) where *c* is log normal with variance 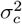. Thus, any trends that depend on environmental noise cannot be directly compared. On the other hand, in accordance with their theory, we do find that increasing neural noise favors canonical olfaction (see *SI Appendix: Fig. S14*). Another important difference is that Lienkaemper *et al*. [47] focus on the setting where the number of neurons *L* is less than the the number of receptors *M*, forcing neurons into coexpression if no receptors are to be neglected. In our setting, *L > M*, which permits neurons to spread out their activity by singly expressing without losing information.

### C. Glomeruli

At the glomerular level, the optimized circuit averages across neurons expressing the same receptor, and uses the transfer function to spread out post-synaptic activity. This has already been understood in terms of efficient coding (see Bhandawat *et al*. [50]), and it is the most intuitive of the three layers. In this way, it serves as a useful sanity check of our computational approach.

Although the simple denoising picture is intuitive, glomerular convergence can also mirror differences in overall levels of receptor abundance (the total number of ORNs which each singly express a given receptor, and the number of copies of that receptor each such ORN expresses). How these differences reflect the environment and experience of the animal has been well-documented experimentally [52, 100], and theoretically analyzed by Teşileanu *et al*. [31]. As it stands, our numerical approach does not support reliable conclusions about optimal allocation of receptor abundance, but this is an area for future development. For the same reason, we did not attempt to make predictions about glomerular convergence in cases of noncanonical expression.

### D. Layer-wise efficient coding

The key structural choice we made in this work was to adopt a layer-wise optimization approach: we first optimized receptor affinities *W* assuming canonical expression *E*, then optimized *E* given fixed *W*, and finally optimized glomerular connectivity *G* given fixed *E* and *W*. Layerwise optimization is both computationally convenient, and—as we will argue below—biologically interpretable.

The risk of such a procedure is that the results could depend on the order of the optimizations. As mentioned before, the analysis of canonical expression represents a chicken-or-egg problem. If we optimize *W* given canonical *E*, then optimize *E* given the resulting *W*, it might be expected that canonical expression would emerge, if this structure is somehow encoded in the optimal *W*. Such a dependence would severely compromise the generality of our results. We therefore sought to check that the finding of generally-canonical expression was robust to the details of the optimization procedure by optimizing expression for different plausible models of the receptor affinity matrix. For these alternative models, optimization resulted in strictly greater levels of canonical expression than the optimized *W*. This suggests that the key takeaway—canonical expression is typically optimal—is not an artifact of our optimization procedure.

As an additional check, jointly optimizing over *W* and *E* yielded similar results to those presented above. The optimized *W* was sparse and close to log-normal, and the optimized *E* did not move far from its noncanonical initialization (see *SI Appendix: Fig. S2*).

From a biological perspective, layer-wise optimization may be a reasonable model for the evolution of the convergent motifs we consider. Organisms are not presented with the opportunity to optimize an entire sensory circuit *ab initio*. Instead, *W, E*, and *G* are tuned by different mechanisms, on different timescales, and in different contexts. We therefore had to approximate the constrained setting in which each motif evolved given limited experimental knowledge.

Chemoreceptor families are evolutionarily ancient and continually evolving [2, 6, 7, 36, 62], so we chose to optimize over *W* first. Since approximate canonical expression is widespread, we plugged in canonical *E* for this optimization. This also permitted direct comparison to the experimental results presented by Si *et al*. [41]. Next, it seems reasonable to ask which gene expression programs are optimal given a broad, mature set of receptors. In the mouse, for example, the relative abundance of olfactory receptors can be adapted over the timescale of just a few hours [52, 101], although this process does not generate coordinated non-canonical expression. This framing also addresses the emergence of different expression programs across organisms which each possess mature receptor families. Hence we optimized over *E* given fixed *W*.

Glomerular convergence, which constitutes an exquisite example of wiring specificity, emerges within the lifespan of the organism by pruning during development [14, 15]. Since examples of highly adaptive connectivity abound in organisms that learn, we chose to study glomerular convergence in the setting of a mature receptor array and largely canonical expression. Hence we optimized over *G* given fixed, optimized *W* and *E*.

These conceptual distinctions are naturally much cleaner than the underlying biology. For example, receptors continue to evolve after a program of noncanonical expression is established [99]. However, our breakdown enables a tractable model and represents a best guess at the constraints which govern the circuit’s evolution. It is of course possible that layer-wise optimization could yield sensitivities and expression patterns that do not lend themselves to glomerular convergence [102]. However, given access to at least as many neurons as receptor types and canonical expression, the simple intuition that glomerular convergence allows denoising suggests that not much is being lost.

One alternative to our approach is an unconstrained end-to-end optimization of the entire circuit. Wang *et al*. [103] recovered largely canonical expression and glomerular convergence using unconstrained optimization on a match-to-prototype classification task. This is qualitatively consistent with our results. Their work, however, differs in assuming a particular classification task, and starts with a very different assumption on the stimulus: they directly model *W* by assuming that the activity of each receptor is independent and uniformly distributed. This differs substantially from the statistics resulting from our optimized sensitivity matrices, and from those measured by Si *et al*. [41]. This kind of unconstrained optimization can shed light on the computation instantiated by the circuit, but cannot capture the path-dependent quirks of biological evolution [104].

Another alternative would be to formalize our context-specific model as a joint dynamical system whose dynamics are decomposed across different timescales. Most simply, this would lead to a setting where we optimize *W* given the expression *E* that is optimized for each sensitivity matrix, *i*.*e*., to study the nested optimization 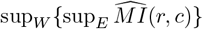. This however could be too flexible—it seems that many organisms have “committed” to canonical expression at the level of gene regulatory mechanisms, for example, and do not have the ability to smoothly modulate receptor coexpression as receptors evolve. Correspondingly, there may be some biological merit to freezing *E* while *W* is optimized.

Fleshing this out completely would require organism-specific knowledge of when and how in evolutionary history each motif emerged. In the absence of such knowledge, we approximated the distinction between the three layers in a biologically plausible way, and treated each one as a separate problem. This piecewise approach limits the normative mathematical claims we can make about the circuit, but it is a closer match to evolutionary dynamics.

### E. Dynamics

Our work does not address the dynamics of the odor landscape and its neural representation in the olfactory bulb. As mentioned above, turbulent transport of air to the sensory epithelium induces high-frequency fluctuations in concentration that are richly informative about identity and location [18, 39]. Here, we have ignored these dynamics in order to focus on a statistical problem of compression: namely, the representation of a high-dimensional chemical space by a finite set of receptors. The temporal challenge of encoding rapidly fluctuating odor signals is equally daunting. Recent work has begun to bridge these statistical and dynamical pictures of olfactory sensing [22, 26, 87, 105–110], but many questions remain regarding how to efficiently encode the temporal statistics of the olfactory world. In this vein, it could be interesting to consider minimal extensions of our model that include temporal filtering [57, 111].

### F. Conclusion

We reiterate in closing that mutual information is only a rough proxy for performance in olfactory processing. For example, organisms may privilege the sensing of rare but critical odorants, even at the expense of more common ones—this would not be captured by our statistical model [65, 69, 112]. Therefore some of our results are unlikely to transfer perfectly to biological circuits. That said, the great variety of olfactory tasks faced by the typical organism suggests mutual information as a reasonable starting point [29, 30, 112].

The correspondence between our results and experimental measurements suggests that these motifs may constitute uniquely accessible solutions to the problem of processing olfactory stimuli. As chemosensation is further explored in non-model organisms [3, 84, 113], the extent to which canonical principles hold will be an interesting point of focus. While receptors always reflect the organism’s chemical environment, the other layers we consider (receptor expression and the downstream routing of receptor activity) are not obviously bound to any stereotypy. Especially interesting will be cases where departures from canonical olfaction are unambiguous, tunable, and clearly advantageous. Since our simplified modeling suggests that canonical olfaction is optimal, such departures should serve as a flag that something interesting is afoot.

## V. METHODS

### A. Generating odor mixtures with tunable statistics

We used a Gaussian copula to generate the binary mixture data *c*_*bin*_ ∈ *{*0, 1} ^*N*^ with tunable mean and covariance. This works as follows.

1. Draw *Z* ∼ 𝒩 (0, Σ) where Σ ∈ ℝ^*N* ×*N*^.
2. Set thresholds *t* = ppf(1 ™ *µ*) where *µ* is the desired mean vector and ppf is the inverse CDF of the standard normal distribution.
3. Set *c*_*bin*_ = **1**_*Z>t*_.

Given Σ_*ii*_ = 1, then *c*_*bin*_ has 𝔼 [*c*_*bin*_] = *µ*. In general, *c*_*bin*_ is not guaranteed to have covariance equal to Σ, but the pairwise covariances 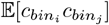 are monotonic, nonlinear functions of Σ_*ij*_.

To model a range of frequencies, with most occurring rarely and some occurring frequently, we drew *µ* itself from a Gamma(*α, λ*) distribution. We parametrized Σ = Σ_*k*_ using a block covariance structure with *k* blocks, as in Figure 1. Each block represented a mixture in the environment emitting correlated odorants. As we varied *k* below *k* = 32 (down to *k* = 4), it was necessary to manually tune *α* and *λ* in order to preserve an approximately fixed odorant frequency distribution while simultaneously achieving the desired correlation structure in the outputs. This is because not all combinations of mean and covariance can be satisfied in binary data.

For low *k*, we also used a “thinning” step in which, after generating the samples, 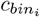 was set to 0 with probability *p*_*k*_. This *ad hoc* intervention was necessary to stabilize the sample-generating algorithm as *k* → 4, ensuring that 𝔼[*c*_*bin*_] was approximately constant across different values of *k* while achieving the desired correlation structure. Thankfully, only minor adjustments to parameters were needed, and exact settings for each *k* are included as parameter files in the codebase so that data can be generated anew as needed. We include plots of the covariance matrix and frequency distribution for each *k* in *SI Appendix: Fig. S17*.

The resulting data *c*_*bin*_ represented binarized odorant mixture samples, with typically 10 (range 1 ™ 100) odorants per sample. We then assigned an *i*.*i*.*d*. log-normal concentration with variance 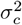 to each odorant that was present in a given sample. This resulted in an odorant vector *c* ∈ ℝ^*N*^ which we passed into our model.

To generate samples with a flat frequency distribution (as in *SI Appendix: Fig. S11*), we repeated the above procedure, but using a constant *µ* = *µ*_0_.

### B. Comparison to other models

The stimulus *c* ∈ ℝ^*N*^ is sensed by the neuron through a panel of *m* receptors. Each receptor has a vector of affinities for each odorant, which is represented by a receptor-by-odorant sensing matrix *W*. Our goal is to describe neural activity as a function of odorant concentration. This obviously involves *W* and *c*, but what form should it take?

One approach is to assume that each neuron expresses just one receptor, and that there is one neuron per receptor type. We will call this the case of single expression, and denote it *r*_*single*_. If the firing rate is simply a linear sum of the (Gaussian) stimulus components

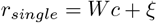

where *ξ* ∼ 𝒩 (0, *σ*_0_) represents neural noise, then *MI*(*r*_*single*_, *c*) = *H*(*r*_*single*_) ™ *H*(*r*_*single*_ | *c*) can be computed analytically.

However, experimental [41, 42, 45, 96] and theoretical [30, 96, 97] works have shown that saturation and other nonlinear effects play an important role in odor coding, especially when the stimulus varies over orders of magnitude. Thus a more realistic activity model is

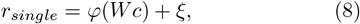

where *φ*(*x*) = 1*/*(1 + *x*^−*n*^) is a Hill function applied element-wise.

We choose a Hill function because experimental work has shown, in the case of monomolecular odorants, that neural activity takes this form as a function of odorant concentration [41], with *n* = 1.46. See Setup in the main text for more details, and *SI Appendix: Fig. S1* for a sweep over Hill coefficients. Note that *φ* is sigmoid when *c* varies over a log scale.

By *r*_*single*_ in (8), we emphasize that this model is making a strong assumption on expression: just one receptor per neuron. Several prior works on efficient coding in olfaction have made this assumption, and within this setup, one can focus on the optimal properties of the sensing matrix *W* (as in [32], [30]) or how neurons should be allocated to each receptor [31]. This assumption is natural, since many organisms exhibit this organization. But here we wanted to target this for study as well, asking how much of canonical olfaction can be recapitulated by efficient coding theory across both layers *W* and *E*. Thus in our model, we include a neuron-by-receptor expression matrix *E* that allows each neuron to express a variable amount of each receptor. This yields the model (1):

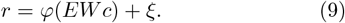

The placement of *E* inside the nonlinearity does *not* constitute a principled mechanistic model integrating receptor signaling. Instead, we are using the Hill function as an empirical model for neural activity in the presence of mixtures.

We use an additive Gaussian model *ξ* ∼ 𝒩 (0, *σ*_0_) for the effect of neural noise. While Poisson noise would more closely model variability in OSN responses [87], differentiating through a Poisson sampler for the purposes of computing gradients was numerically challenging (requiring a continuous relaxation). For simplicity, we therefore used Gaussian noise. For all analyses except the sweep over neural noise, we set the magnitude of the noise to be *σ*_0_ = 0.1. This was to match the approximate mean activity 𝔼[*r*] when computed over samples—see Glomeruli and *SI Appendix: Fig. S16* for histograms of activity.

### C. Details of mutual information maximization

We leverage a variational formulation of mutual information first proposed by Nowozin *et al*. [61]. Following their presentation, we give a short overview of the theory underlying this approach, which was first presented by Nguyen *et al*. [114]. The *f-divergence* between two distributions *P* and *Q* defined on a domain with 𝒳 densities *p*(*x*) and *q*(*x*) is defined as

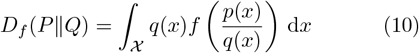

The mutual information *MI*(*g, c*) is a special case of this: namely the Kullback-Leibler divergence between the joint distribution *p*(*x*) = *p*(*g, c*) and the product of marginals *q*(*x*) = *p*(*g*)*p*(*c*), with *f* (*u*) = *u* log(*u*).

If *f* is convex and lower semi-continuous, then we can write the Fenchel conjugate

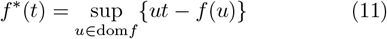

The functions *f* and *f* ^∗^ are dual in the sense that *f* = *f* ^∗∗^. Then we can write

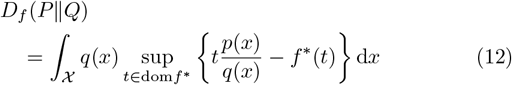

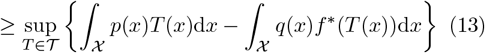

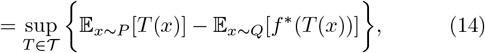

where 𝒯 is an appropriately-chosen class of test functions *T* : 𝒳 → ℝ. We have samples from the joint distribution *p*(*g, c*), and we obtain samples from the distribution *p*(*g*)*p*(*c*) by shuffling activity. Thus, we can estimate the expectations in (14).

We then simply parameterize *T* using a small neural network, and optimize jointly over the layer in question and the parameters of *T*. Tschannen *et al*. [115] gave a thorough discussion of critic architectures. Following their recommendations, we used an inner product architecture for *T* so that *T* (*g, c*) = *χ*(*g*)^⊤^*ψ*(*c*), where *χ* and *ψ* are separate two-hidden-layer networks with width = 128. To sanity-check that our results did not depend strongly on the choice of critic network, we performed spot-checks using other critic architectures suggested by Tschannen *et al*. [115].

Such approaches are not without controversy. McAllester and Stratos [58] have argued that, for the Kullback-Leibler divergence (KLD), the expectations in (14) cannot be estimated without exponentially many samples. This is ultimately due to the fact that the KLD has *f* ^∗^(*t*) = exp(*t ™* 1). This theoretical result prompted investigation into why such estimators are still useful in applications such as representation learning [115].

Here, we sidestep this problem by using the Jensen-Shannon divergence (JSD), a symmetrized version of the Kullback-Leibler divergence (KLD) which has *f* ^∗^(*t*) = ™ log(2 ™ exp(*t*)), thus avoiding the need for exponentially many samples. The greater stability and robustness of this Jensen-Shannon-based estimator compared to the exact KL formulation were empirically established by [59] and [60]. These variational approaches have gained considerable popularity in the machine learning community, to the point that maximizing mutual information is no longer considered an intractable problem [56, 59, 60].

### D. Constrained optimization using mirror descent

When optimizing over *W, E*, and *G*, we must account for the constraints on each matrix: *W* must have positive elements, and both *E* and *G* must have non-negative elements and have all row sums equal to unity (*i*.*e*., each row lies in a simplex). To handle these constraints, we use the framework of mirror descent [116, 117]. For a pedagogical introduction to mirror descent, we direct the reader to Vlad Niculae’s blog post [118].

This approach requires specifying a *ψ* function which maps from the primal (constrained) space to the dual (unconstrained space), and *ϕ* function which maps from the dual space back to the primal space. So for *W*, to ensure positivity, we have used *ϕ*(*u*) = exp(*u*) *>* 0. For *E* and *G* optimizations, we used *ϕ*(*u*) = softmax(*u*)_*i*_ = exp(*u*_*i*_)*/* ∑_*i*_ exp(*u*_*i*_). Mirror descent is equivalent to using “natural” gradient descent [119] in the dual space, but enjoys favorable numerical properties [117]. It greatly increased the stability of our optimization procedure.

All gradients were computed via automatic differentiation using the JAX library [120]. Initializations were as discussed in the main text: for *W*, we initialized using a scaled log-normal. For *E*, we varied the initialization depending on the analysis. In Figure 3, we initialized to random expression. In *SI Appendix: Fig. S9*, we initialized to canonical expression. Otherwise, we initialized to noncanonical expression. Typically this meant 3-7 receptors per neuron with roughly equal expression, as in Figure 5. However, in *SI Appendix: Fig. S13*, we initialized to 1-5 receptors per neuron with roughly equal expression. This was to respect the block structure of *W* when modeling coexpression of correlated receptors. For *G*, we initialized to random connectivity (Figure 6).

## ACKNOWLEDGMENTS

We are grateful to A. D. T. Samuel and all the authors of Si *et al*. [41] for making their data publicly available. J.C.F.C. thanks D. M. Zimmerman and I. Chandok for helpful discussions regarding the evolution and biophysics of receptors. We thank C. Lienkaemper, M. A. Younger, and G. K. Ocker for inspiring discussions regarding non-canonical expression. Moreover, we are indebted to A. D. T. Samuel and the members of his group, and to G. K. Ocker and M. A. Younger, for their thoughtful comments, which have helped us improve our manuscript.

J.C.F.C. was supported by the Harvard Biophysics Graduate Program through training grant T32GM158477 from the National Institutes of Health. F.P. was supported by the Harvard Center for Brain Science (CBS)-NTT Fellowship Program on the Physics of Intelligence. V.N.M. was supported by grant number A47994 from NTT Research, Inc. J.A.Z.-V. was supported by the Office of the Director of the National Institutes of Health under Award Number DP5OD037354, and by a Junior Fellowship from the Harvard Society of Fellows. This work has been made possible in part by a gift from the Chan Zuckerberg Initiative Foundation to establish the Kempner Institute for the Study of Natural and Artificial Intelligence at Harvard University.

## AUTHOR CONTRIBUTIONS

J.C.F.C., V.N.M., and J.A.Z.-V. conceptualized the project, with input from F.P. J.C.F.C. wrote code, performed simulations, prepared figures, and wrote an initial draft of the paper, with input and assistance from all authors. All authors contributed to editing the manuscript. J.A.Z.-V. and V.N.M. jointly supervised this project.

## DATA AND CODE AVAILABILITY

Code to perform all optimizations and reproduce all figures is available on GitHub at https://github.com/VNMurthyLab/olfactory-ec.

The dataset of estimated *Drosophila* larva receptor affinities from Si *et al*. [41] that we analyzed in Figure 2 was downloaded from A. D. T. Samuel’s lab GitHub, where it is available under an MIT License: https://github.com/samuellab/Larval-ORN/blob/master/Figure3/results/MLEFit.mat.

## Supporting Information

**Appendix A: Estimating the minimum effective sensitivity**

In our analysis, with the exception of Figure 2, we have initialized the sensitivities *W*_*ij*_ to the scale 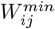 below which *W*_*ij*_ is effectively 0. This allows us to confirm, for example, that broad receptor tuning is not an artifact of initialization (see *SI Appendix: Figs. S3, S7*). Knowing 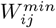 is also necessary to understand the range of biologically meaningful tuning, which differs from the range of the optimized *W*_*ij*_. For example, many of the optimized elements *W*_*ij*_ are far below working machine precision (∼ 10^−16^), but this does not mean that the optimal *W* spans more than 16 orders of magnitude. Even biologically, for example, *W*_*ij*_ = 10^−14^ is equivalent to *W*_*ij*_ = 0, since none of our odor concentrations are close to 10^14^. To correctly interpret our results, we must understand the minimum effective value for *W*_*ij*_. We discuss below how that minimum relevant value is set by the maximum concentration *c*_*j*_.

Assuming canonical expression for simplicity, the mean activity of a neuron expressing the *i*-th receptor is given by

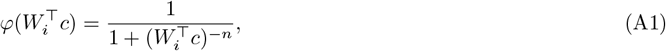

where *W*_*i*_ is the vector of sensitivities for the *i*-th receptor. To determine the minimum relevant sensitivity, we would like to estimate the smallest *W*_*ij*_ such that changing the concentration *c*_*j*_ produces a detectable change in 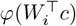. We do so by freezing all components of the concentration vector but the *j*-th, and write

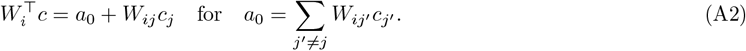

Then, we linearize in *W*_*ij*_*c*_*j*_ around the half-occupancy point *a*_0_ = 1, which is where the Hill function is maximally sensitive to changes in inputs:

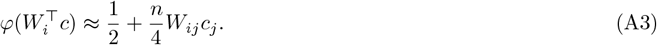

In the absence of neural noise, any contribution above working precision results in an numerically meaningful change in the activity. But for non-zero neural noise levels *σ*_0_, we must estimate a minimum distinguishable difference. For example, in our modeling of the adult fly, *σ*_0_ = 0.1, and we have approximately 21 neurons per receptor. So the effective noise is approximately 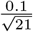. Therefore, a rough estimate for a meaningful difference in mean activity compared to the noise is to take *W*_*ij*_*c*_*j*_ to scale with the standard deviation of the effective noise, *i*.*e*.,

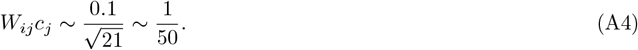

Other choices for this value simply result in a different prefactor in the final expression for 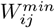. To obtain the lowest possible 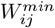, we set *c*_*j*_ = *c*_*max*_. This results in

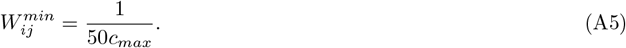

This reasoning ensures that 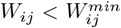 is functionally equivalent to *W*_*ij*_ = 0. In other words, the spread of *W*_*ij*_ below 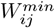 is not biologically meaningful. This allows us to make reasonable claims about the distribution of *W*_*ij*_, as in Fig. 2. Finally, recall that *c* is log-normal, so the maximum *c*_*j*_ is on the order of *c*_*max*_ = exp(*µ* + 3*σ*). In this way, the value of 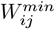 (and thus, the definition of the sparsity of *W*) depends on the environmental noise.

This argument is purely heuristic, and only intended to guide our interpretation of numerical results. We do not use it as the basis for any biological claims. In particular, we have not conducted any analyses that depend on a precise estimate of 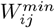. For example, Qin *et al*. [30] report that as *σ*_*c*_ increases, *W*_*opt*_ becomes less sparse (more entries are non-zero). This is plausible, but difficult to confirm in our hands because the meaningful threshold for sparsity is also changing with *σ*_*c*_. Furthermore, one could take *c*_*max*_ = exp(*µ* + 3*σ*), but one might take a more typical value (exp(*µ* + 2*σ*)) or a more extreme one (exp(*µ* + 4*σ*)). Each choice for *c*_*max*_ results in a different 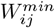.

---

**FIG. S1.**
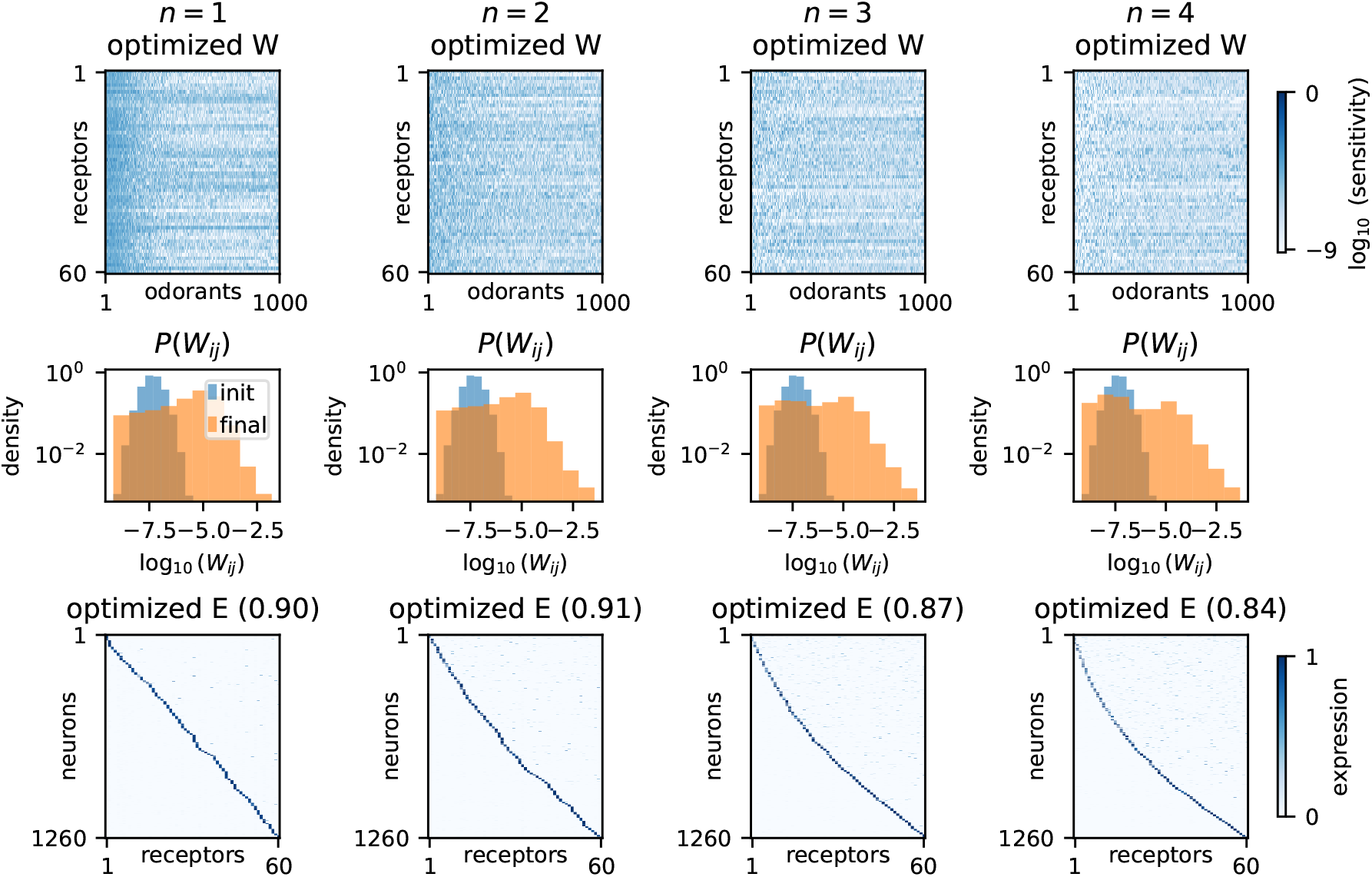
Effect of the Hill coefficient *n* on optimized *W* and *E. E* optimizations are performed using shuffled *W*_*opt*_. In the histograms, blue represents the initial *W*_*ij*_, and orange the optimized *W*_*ij*_. *E* was initialized to noncanonical olfaction as in Figure 5a, b. The “canonical score” (as defined in the caption of Figure 5) is shown in parentheses for each optimized *E* matrix. A score of 1 indicates perfect single receptor expression in each neuron. Environmental parameters were set to *σ*_*c*_ = 2 and 64 sources.

**FIG. S2.**
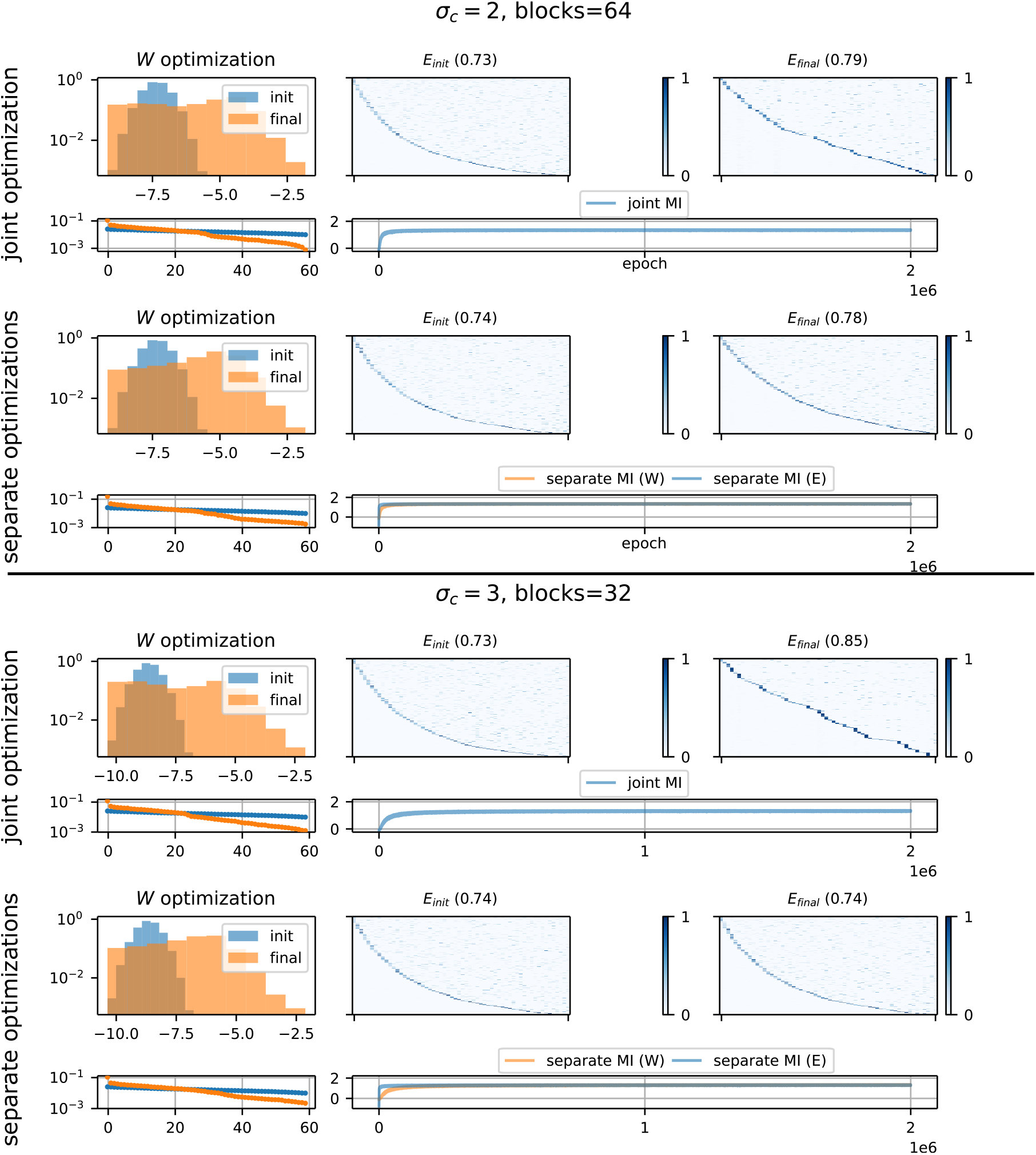
Comparing the joint optimization over *E* and *W* (rows 1 and 3) to layer-wise optimization (first *W* then *E*, rows 2 and 4) for two different realistic settings of environmental parameters. In the first column, the distribution of the non-zero entries of *W*_*ij*_ is shown on a log_10_ scale. Below the histograms are the respective spectra of *W*_*init*_ and *W*_*opt*_. The same low-rank structure is obtained for *W* in both the joint and layer-wise cases (see Fig. S10 for sweep over parameters). In the right two columns, *E*_*init*_ and *E*_*final*_ are shown, and trajectories of the optimizations are shown below. *E* moves little from optimization, a result of the low rank structure in *W*. The joint results are consistent with those obtained by layer-wise optimization, as presented in the main text.

**FIG. S3.**
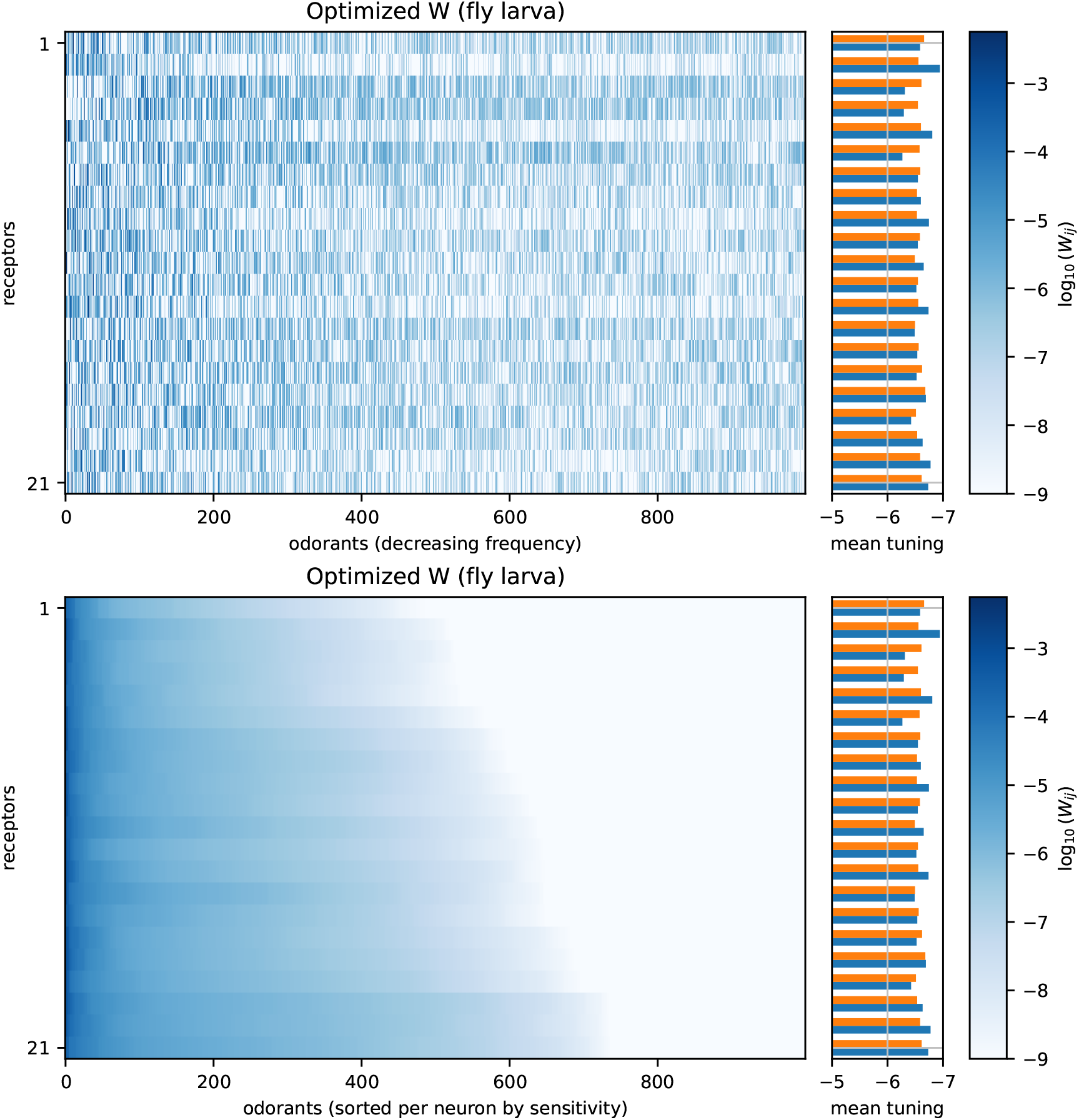
Receptor tuning curves in the optimized *W* matrix for the fly larva. The same data is shown twice: for the top panel, odorants are sorted by frequency, whereas for the bottom panel, odorants are sorted within each receptor by sensitivity. Bars on the right indicate the mean log-sensitivity of each receptor (blue is true, orange is shuffled). The receptor mean tuning is more variable than would be expected by chance (blue bars vary more than orange bars). This is consistent with work [11] showing that some receptors are very narrowly tuned (or almost exclusively inhibitory), while others are broadly tuned. Environmental parameters were set to *σ*_*c*_ = 3 and 32 sources.

**FIG. S4.**
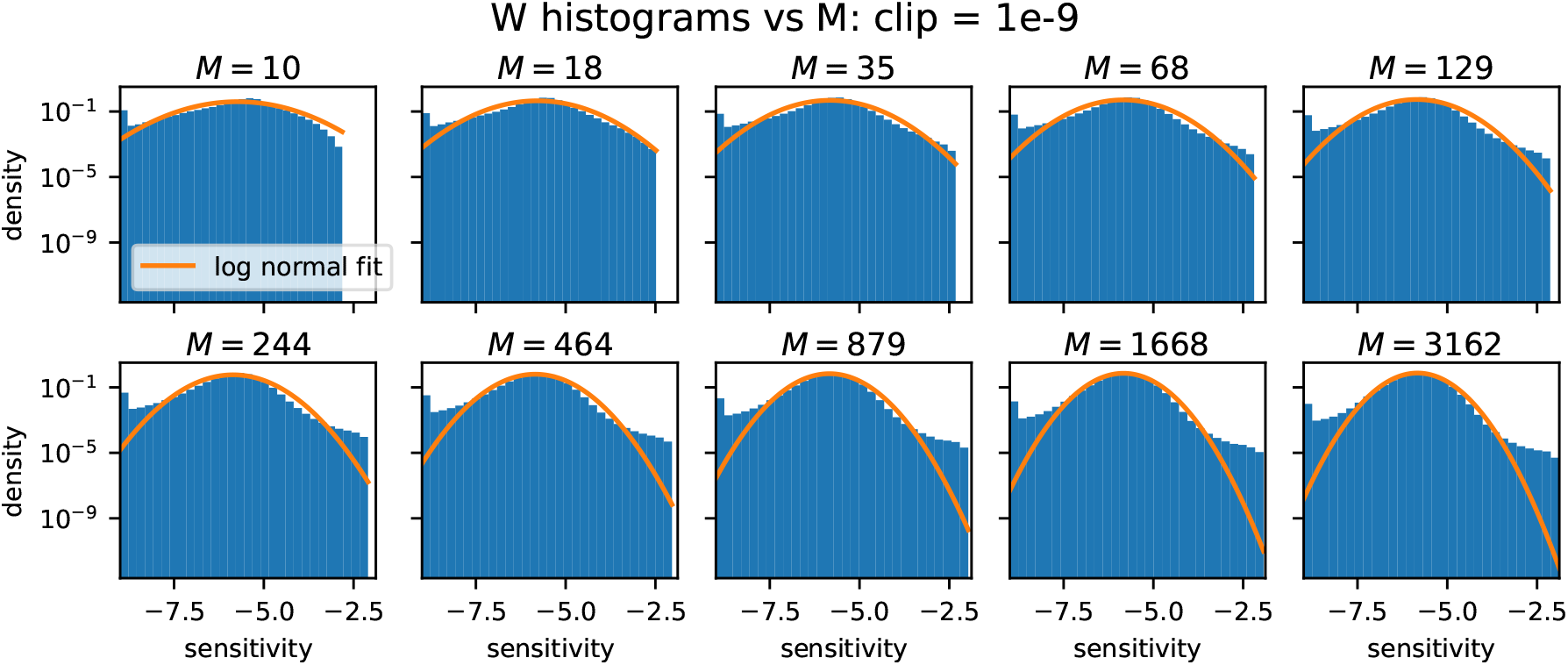
Histograms of *p*(log_10_(*W*_*ij*_)) after optimization, and log-normal fits. Note that the log density is plotted. The heavy right tails of the true distribution compared to the log-normal drive the emergence of specialists for large *M* (second row). Environmental parameters were set to *σ*_*c*_ = 3 and 32 sources.

**FIG. S5.**
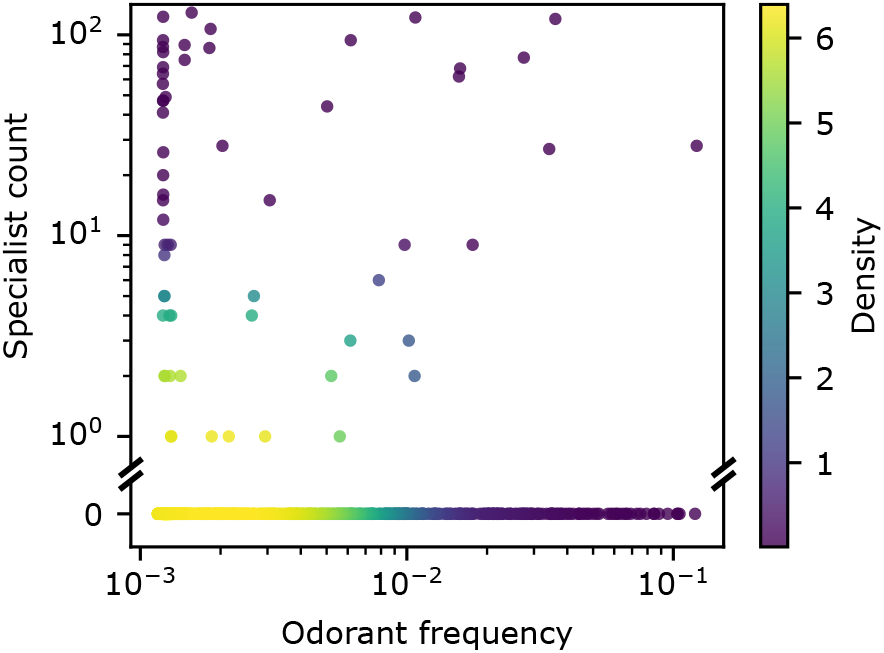
Specialist receptor count vs. odorant frequency. “Specialist count” is the number of times the odorant was the target ligand of a specialist receptor in any of the 200 simulations shown in Figure 2e (20 each for 10 different values of *M*). Since most odorants were never targeted by specialists, we were not powered to analyze trends for given values of *M*. Environmental parameters were set to *σ*_*c*_ = 3 and 32 sources.

**FIG. S6.**
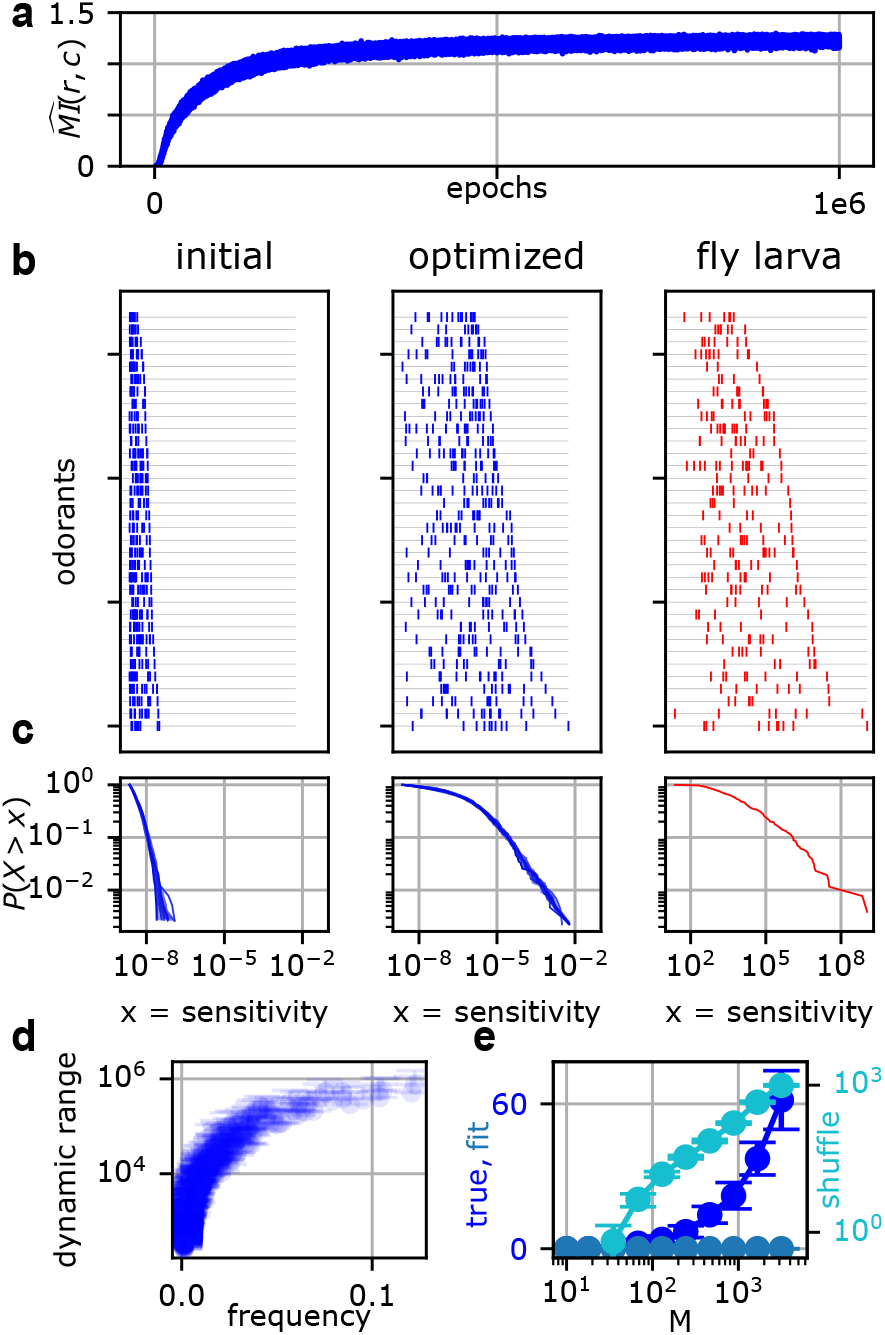
Optimized *W* distribution under initialization to minimum *W*_*ij*_ (compare to Figure 2 in main text.) The only significant difference is in the “shuffle” curve for panel e, which is shifted dramatically upward (note log scale). This reflects the fact that, for large *M*, most sensitivities do not need to move under optimization. When *W* is shuffled, this causes many specialists to emerge. However, in the true *W*, the number of specialists was comparable to Figure 2e (mean 62 specialists for *M* = 3162 receptors rather than mean 30 specialists for the 1*/E*[ ||*c*||] initialization in the main text). This indicates some pressure against developing too many specialists, even in an over-parameterized (*M > N*) regime. See Appendix *A* for a discussion of how the minimum effective *W*_*ij*_ is computed. Environmental parameters were set to *σ*_*c*_ = 3 and 32 sources.

**FIG. S7.**
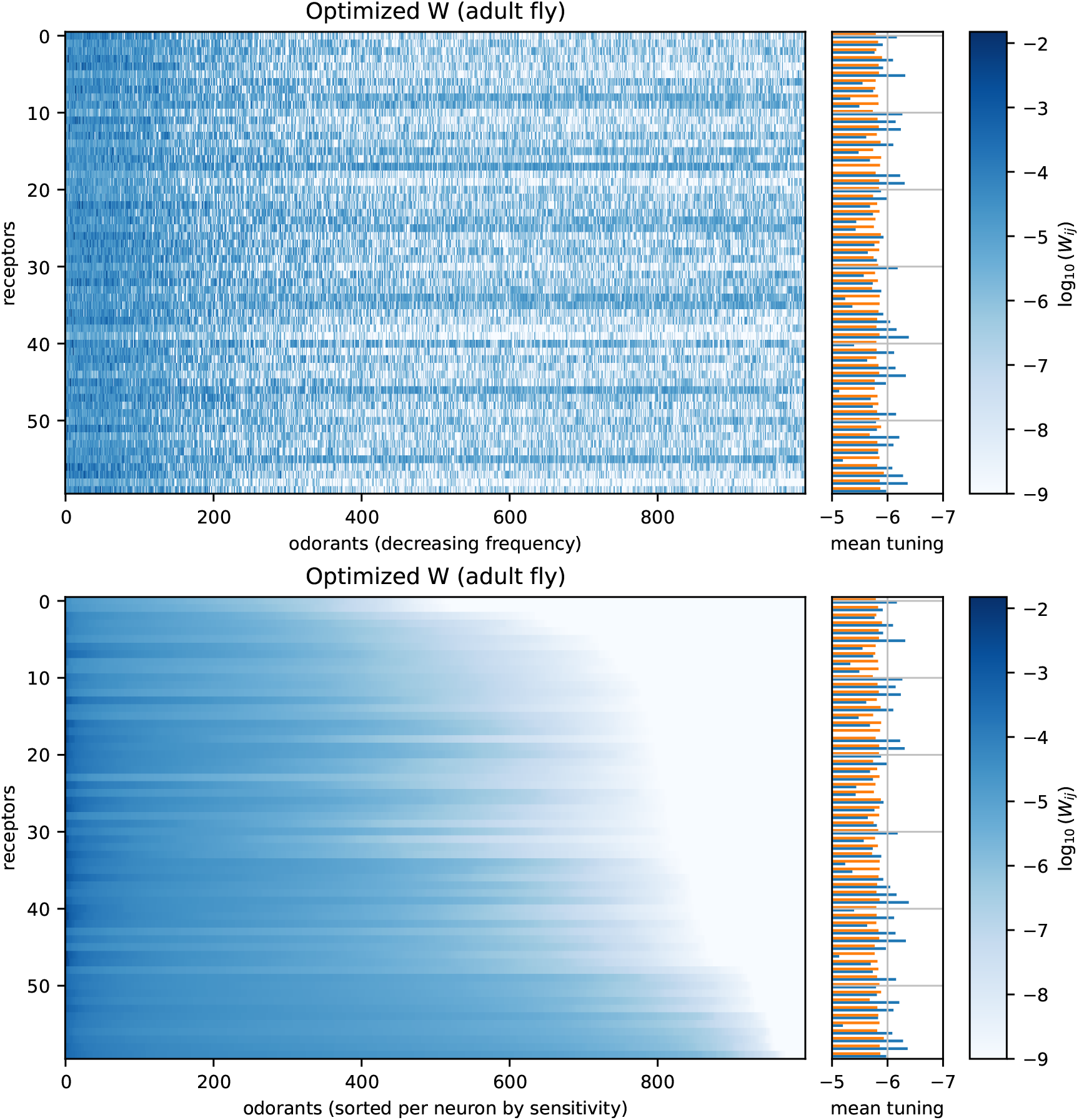
Receptor tuning curves in the optimized *W* matrix for the adult fly. The same data is shown twice: for the top panel, odorants are sorted by frequency, whereas for the bottom panel, odorants are sorted within each receptor by sensitivity. Bars on the right indicate the mean log-sensitivity of each receptor (blue is true, orange is shuffled). The receptor mean tuning is more variable than would be expected by chance (blue bars vary more than orange bars). This is consistent with work [11] showing that some receptors are very narrowly tuned (or almost exclusively inhibitory), while others are broadly tuned. Environmental parameters were set to *σ*_*c*_ = 2 and 64 sources.

**FIG. S8.**
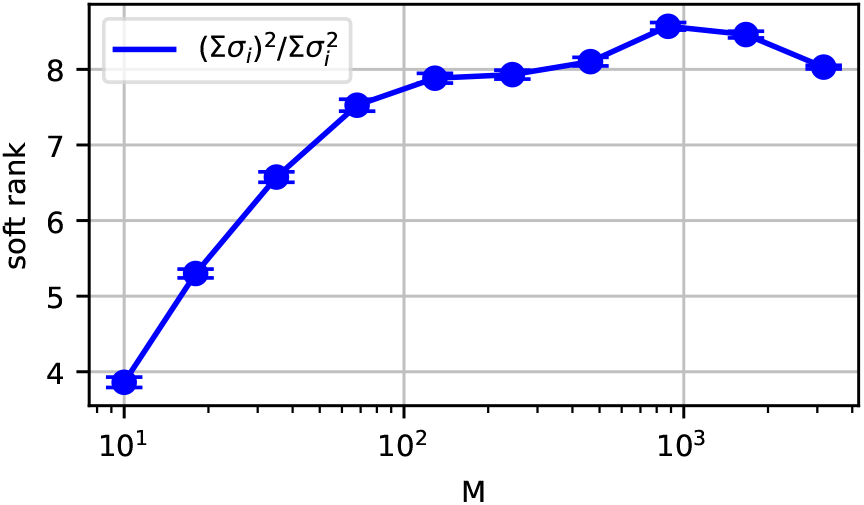
Soft rank of *W* vs. receptor count *M*. The soft rank is computed using 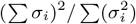, where *σ*_*i*_ are the singular values of *W*. This measure is 1 when *W* is rank 1 and *M* when *W* is full-rank. Error bars indicate the standard deviation over 20 runs. Environmental parameters were set to *σ*_*c*_ = 3 and 32 sources.

**FIG. S9.**
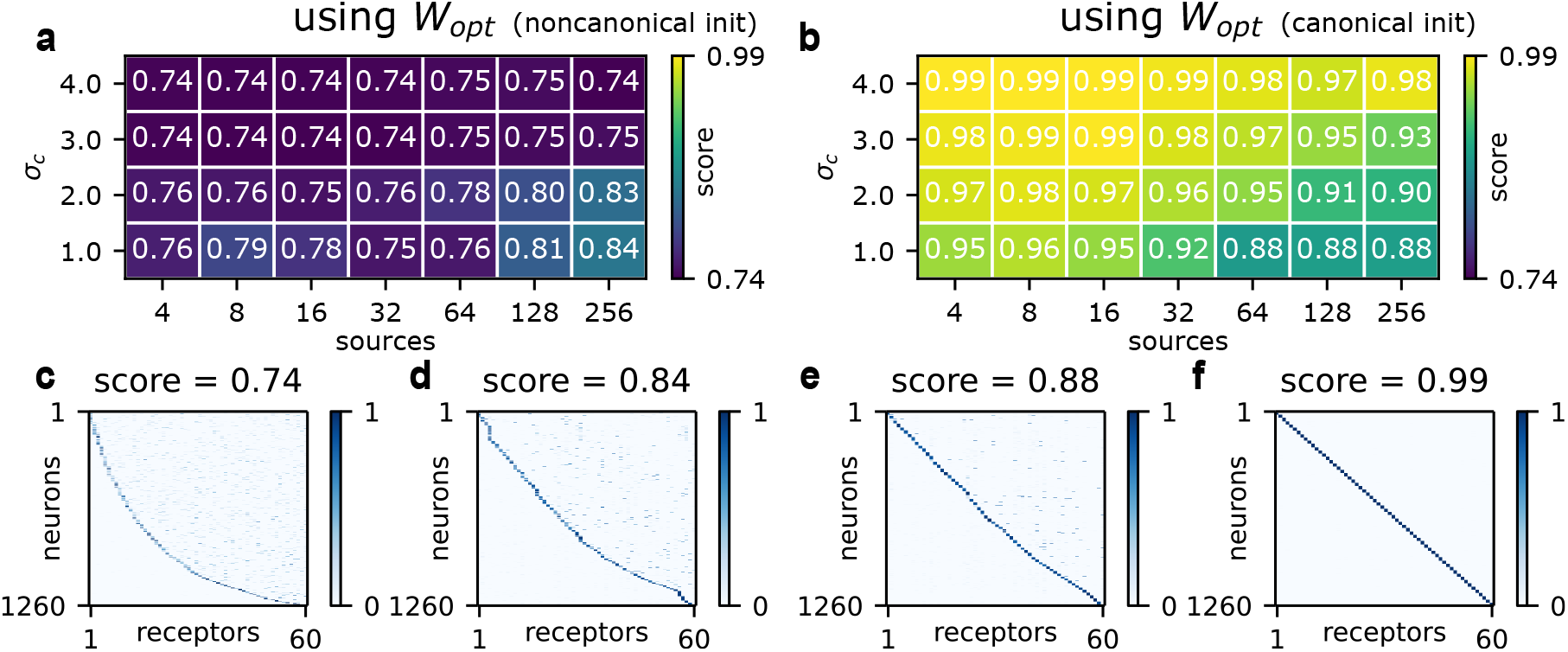
Phase diagram of *E* optimization given *W*_*opt*_, comparing canonical and noncanonical initialization. Only the optimizations in the bottom right corner (low noise, many sources) of the phase diagram move meaningfully from initialization.

**FIG. S10.**
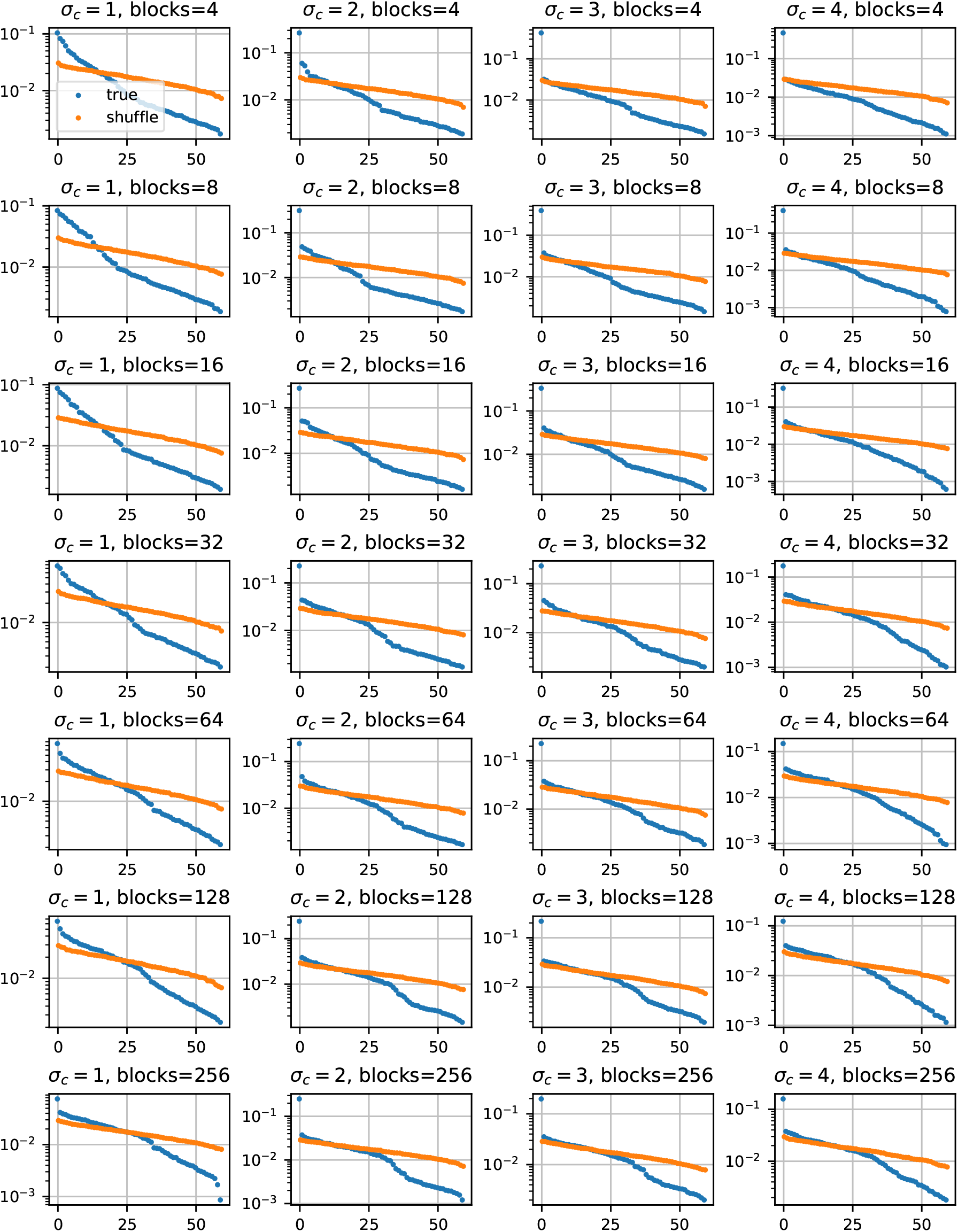
Eigenvalue spectra of optimized *W* across environments, given the standard odorant model (most odorants are rare, some are common, as in Figure 1).

**FIG. S11.**
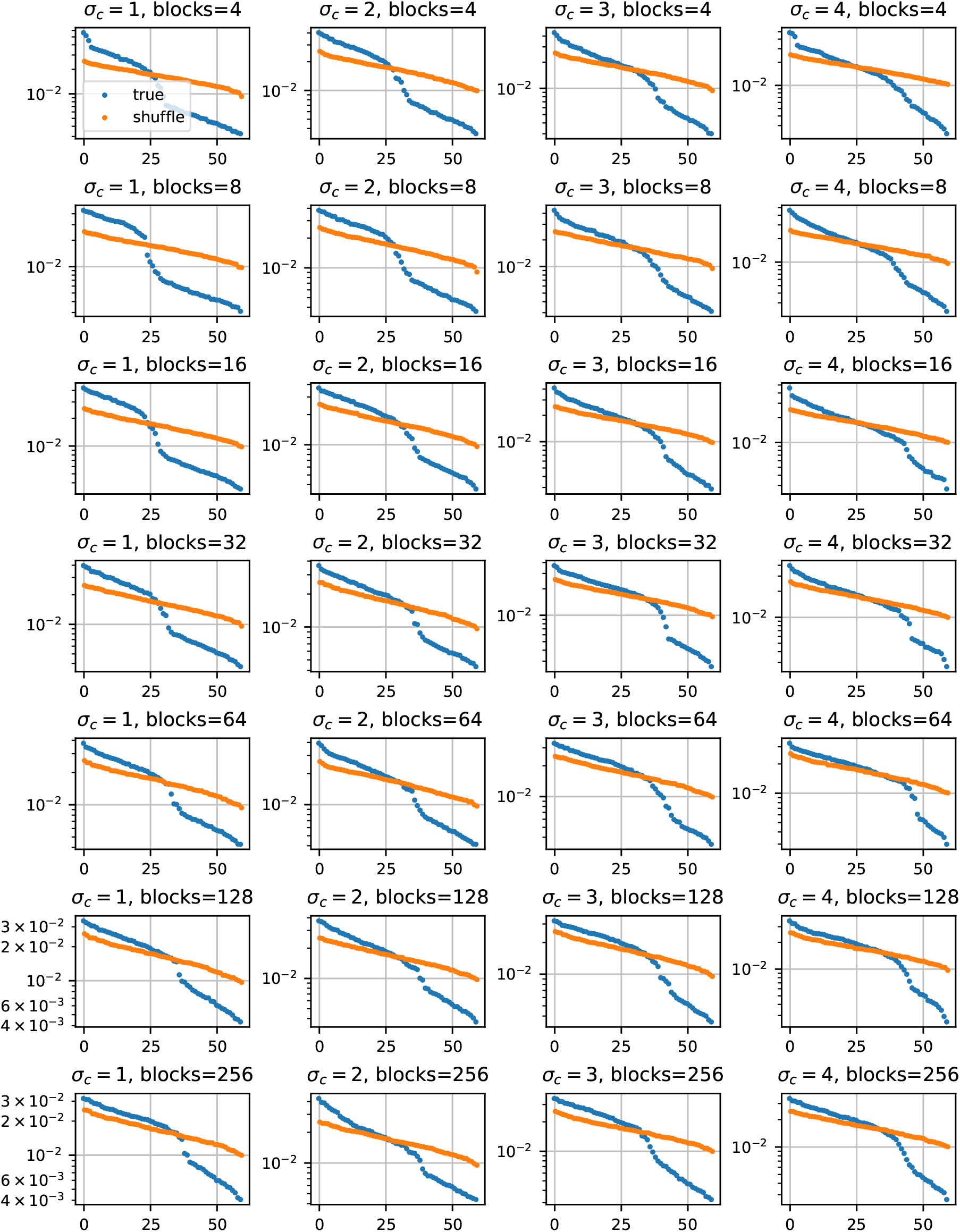
Eigenvalue spectra of *W* under an odorant model with constant frequencies (so all *N* = 1000 odorants occur with equal probability). The low rank structure in Fig. S10 is abolished.

**FIG. S12.**
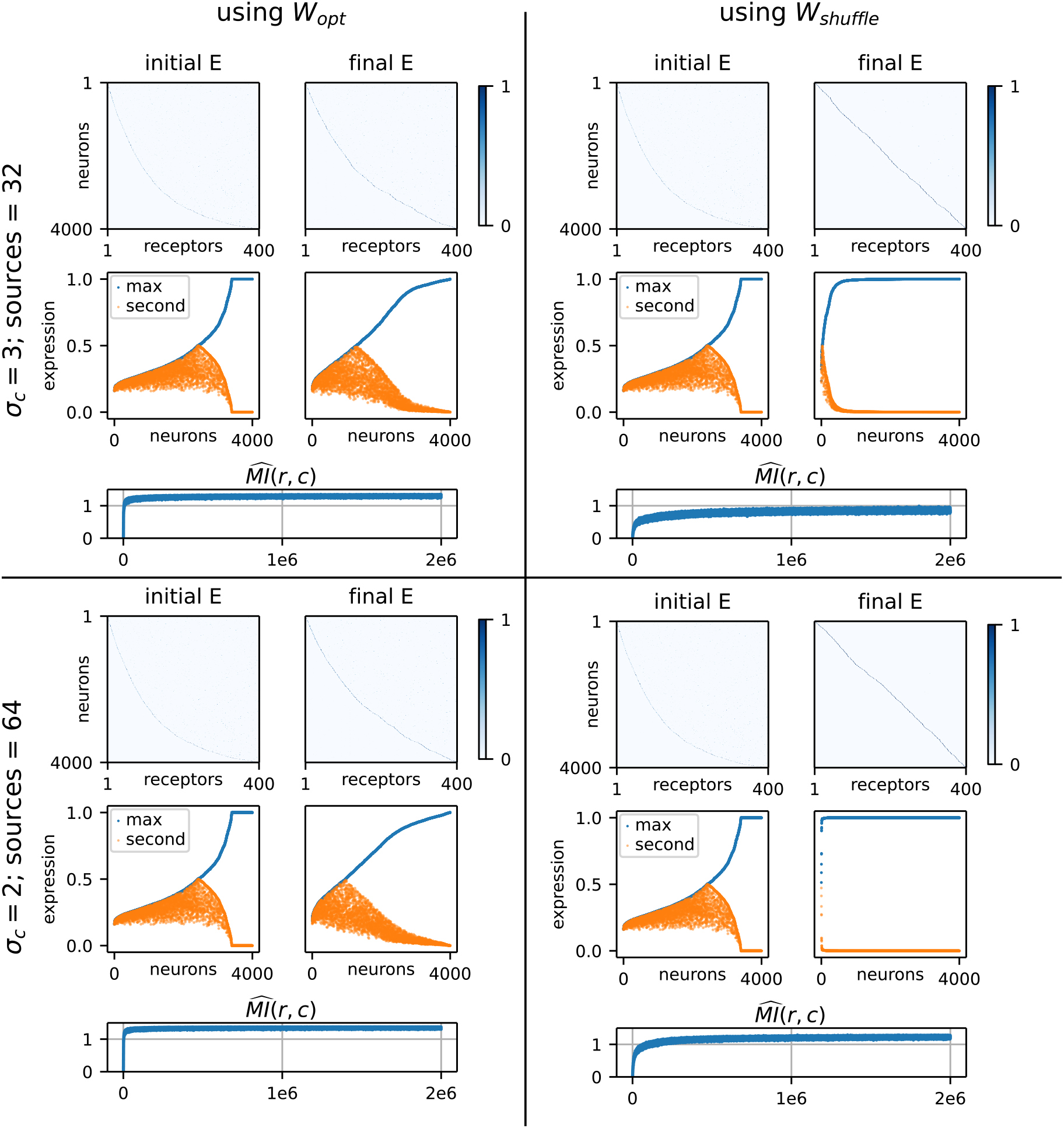
Patterns of optimal expression for two different *W* matrices (*W*_*opt*_ and *W*_*shuffle*_, columns) at two different points in environmental parameter space (rows). At this system size, the heatmaps are harder to interpret, so below each expression plot we visualize expression by plotting the maximum entry in each row (blue) and the second-to-max entry (orange). Similar to the adult fly, we find that *W*_*opt*_ permits noncanonical expression, but *W*_*shuffle*_ encourages precise one neuron-one receptor expression. Note that, for this analysis, *N* = 10000 odorants were used, rather than *N* = 1000. This was to maintain a meaningful bottleneck

**FIG. S13.**
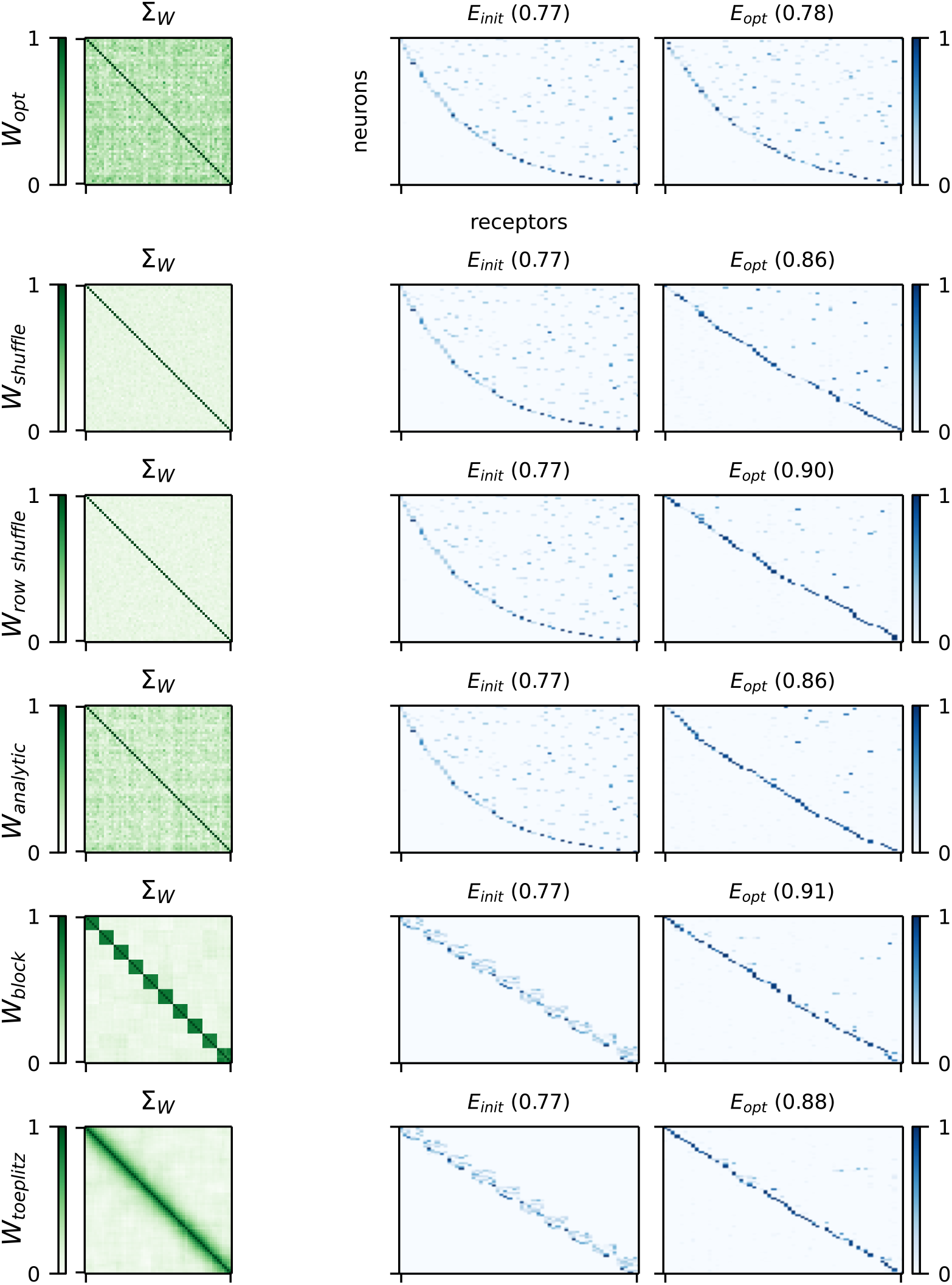
Optimized expression given a variety of different *W* matrices. Environmental parameters are set to 64 sources and *σ*_*c*_ = 2. For *W*_*block*_ and *W*_*toeplitz*_, the E matrices were initialized to exhibit coexpression of correlated receptors (coexpression by descent) [51]. This did not lead to more noncanonical expression than the shuffled *W*, in which unrelated receptors were coexpressed. *W*_*analytic*_ is generated by fitting a log normal to *W*_*opt*_, then masking by the same sparsity structure present in *W*_*opt*_. This induces a non-negligible correlation structure in *W*_*analytic*_.

**FIG. S14.**
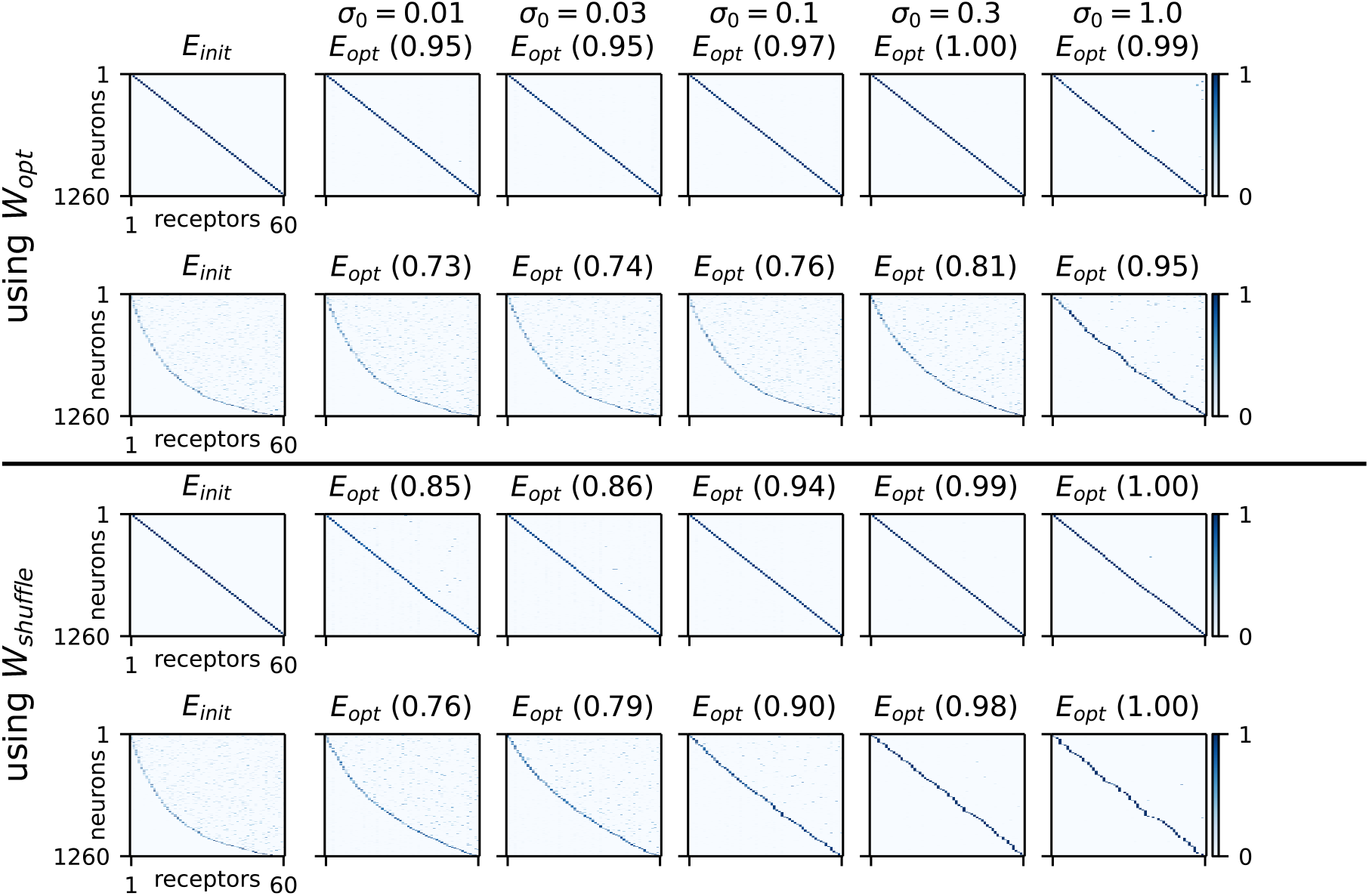
Sweeping across neural noise *σ*_0_. The canonical score (as defined in Figure 5) is shown in parentheses. Both canonical (first and third rows) and noncanonical (second and fourth rows) initializations are shown. For very low levels of neural noise, some noncanonical expression is tolerated. For *σ*_0_ ∈ [0.1, 1.0], canonical olfaction is strongly favored. Environmental parameters are set to 64 sources and *σ*_*c*_ = 2.

**FIG. S15.**
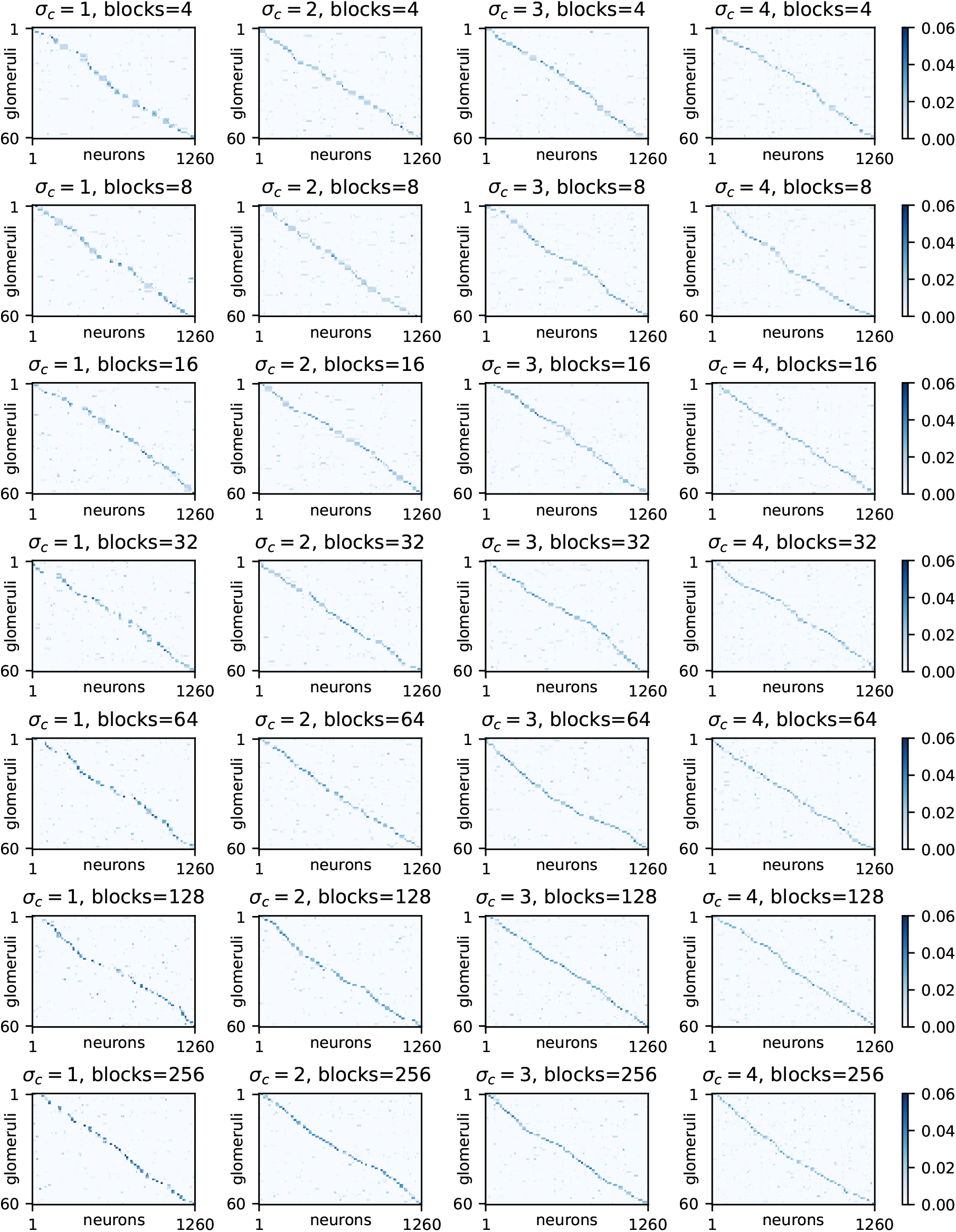
Glomerular convergence across environmental parameters using shuffled *W* and corresponding optimized *E*.

**FIG. S16.**
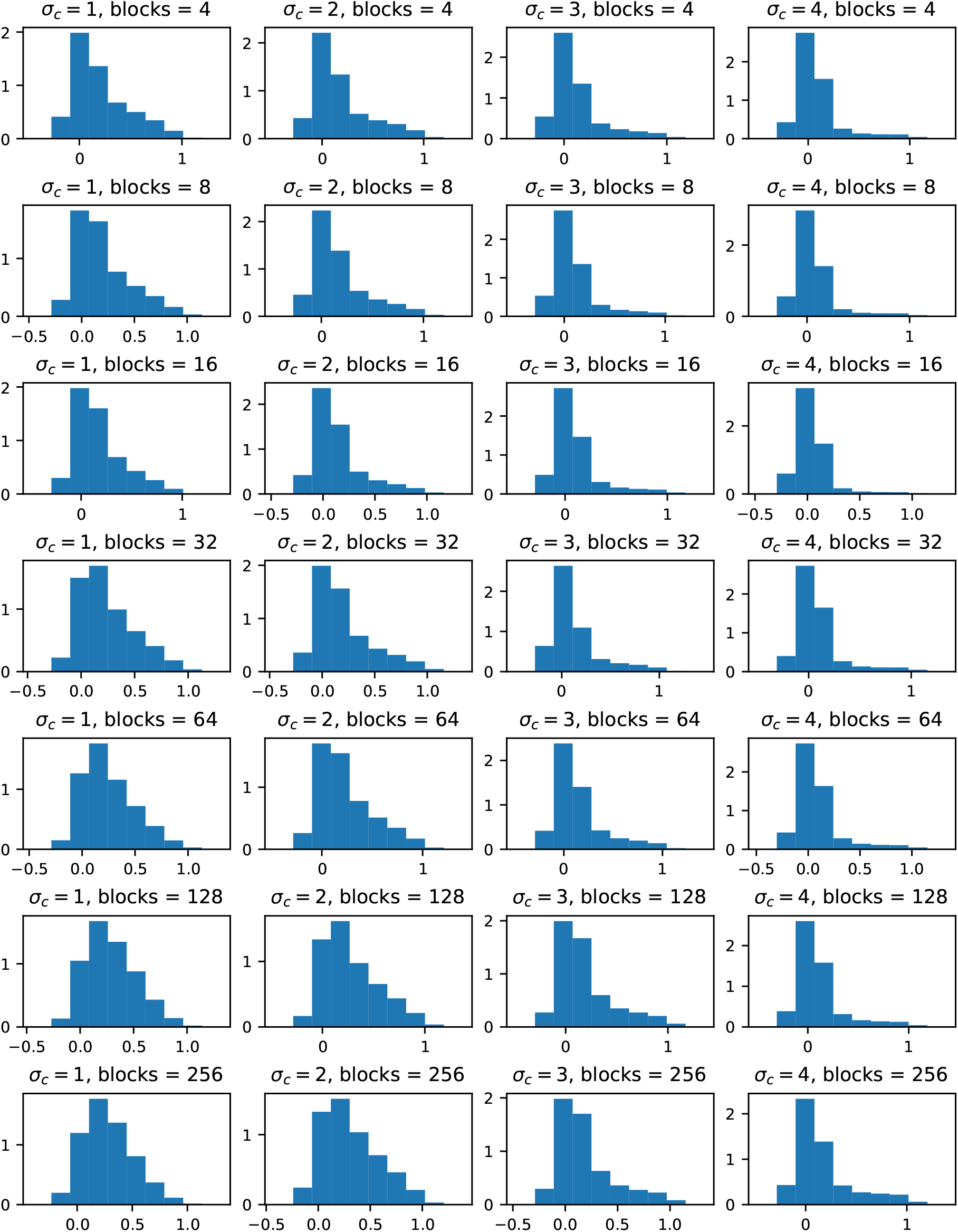
ORN activity distributions across environmental parameters using shuffled *W* and corresponding optimized *E*.

**FIG. S17.**
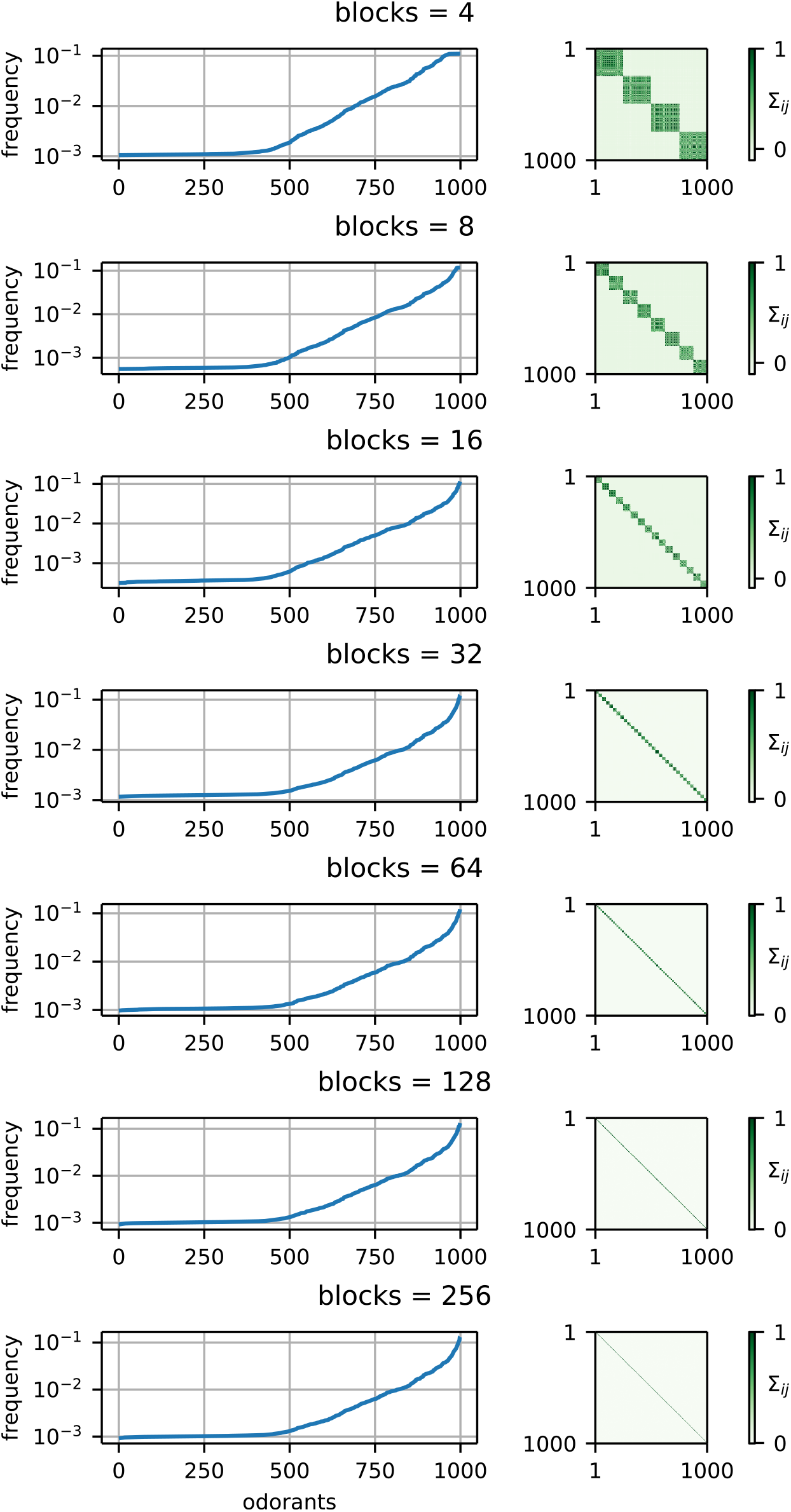
Mean and covariance of the binarized odorant vector *c*_*bin*_ as the number of blocks *k* is varied.

